# Human brain changes after first psilocybin use

**DOI:** 10.1101/2024.10.11.617955

**Authors:** T Lyons, M Spriggs, L Kerkelä, FE Rosas, L Roseman, PAM Mediano, C Timmermann, L Oestreich, BA Pagni, RJ Zeifman, A Hampshire, W Trender, HM Douglass, M Girn, K Godfrey, H Kettner, F Sharif, L Espasiano, A Gazzaley, MB Wall, D Erritzoe, DJ Nutt, RL Carhart-Harris

## Abstract

Psychedelics have robust effects on acute brain function and long-term behavior but whether they also cause enduring functional and anatomical brain changes is unknown. In a placebo-controlled, within-subjects, electroencephalography, and magnetic resonance imaging study in 28 healthy, entirely psychedelic-naive participants, anatomical and functional brain changes were detected from one-hour to one-month after a single high-dose (25 mg) of psilocybin. Increases in cognitive flexibility, psychological insight, and well-being were seen at one-month. Diffusion imaging done before and one-month after 25mg psilocybin revealed decreased axial diffusivity bilaterally in prefrontal-subcortical tracts that correlated with decreased brain network modularity over the same time period. Decreased modularity also correlated with improved well-being. Increased cortical signal entropy at 1– and 2-hours post-dosing predicted improved psychological well-being at one-month. Next-day psychological insight mediated the entropy to well-being relationship. All effects were exclusive to 25mg psilocybin; no effects occurred with a 1mg psilocybin ‘placebo’ dose.

## INTRODUCTION

Psilocybin (4-PO-HO-DMT) is the precursor of psilocin (4-HO-DMT), a serotonin receptor agonist. Converging evidence supports a role for serotonin 2A receptor (5-HT2AR) agonism in eliciting the characteristic brain and behavioural effects of this and related psychedelics in humans ^1,2^. Pre-clinical research supports an association between 5-HT2AR agonism and increased ‘neuroplasticity’ ^3–7^, although see ^8^.

Clinical trials have assessed psilocybin-therapy for psychopathology ^9–14^. Human research has found enduring psychological changes after single high-dose psilocybin ^15^, including increased cognitive flexibility in depressed patients ^16^ (although see ^17^ with LSD) and improved well-being in diverse samples ^18,19^. Several human neuroimaging studies have observed changes in acute brain function with psychedelics ^20–24^. Long-term brain changes are less well characterized, but see ^25,26^ in healthy volunteers and ^16,27–30^ in depression, plus Table S1 for a list of relevant studies.

In the present work, we sought to address important knowledge gaps regarding human brain changes with psilocybin. Electroencephalography (EEG), functional magnetic resonance imaging (fMRI) and diffusion tensor imaging (DTI) recordings were done in healthy human volunteers receiving their first-ever high-dose of a psychedelic (25mg psilocybin). Brain (fMRI/DTI) and behavioural outcomes were assessed at baseline and one-month post-dosing. EEG was done during dosing. Due to prior evidence of enduring behavioural changes after psilocybin ^16,26,27,29^, a fixed-order, repeated measures design was used; participants received an initial ‘placebo’ dose of psilocybin (1mg) one-month prior to a subsequent 25mg dose. Dosing was oral. See Figures S2-3 for study design details.

## RESULTS

### Demographics

Twenty-eight healthy volunteers participated, mean age = 41 (SD=8.7), 43% female. See Figure S4 for a demographics table.

### Neuroimaging outcomes

#### Acute brain effects

##### Time-dependent changes in LZc and spectral power (EEG)

Eyes-closed, task-free ‘resting-state’ brain activity during each of the two psilocybin dosing sessions (i.e., 1mg and 25mg, one-month apart) was investigated using EEG performed at pre-dose baseline and 1-, 2-, and 4.5-hours post-dosing. Robust pre vs acute (i.e., ‘on-drug’) brain changes were observed after the 25mg dose, as revealed by *Dose* (1mg vs 25mg) x *Time* (0-4.5h) linear mixed-effects modelling. There were significant (*p*<0.001) increases in the informational entropy of spontaneous scalp potentials (Lempel-Ziv complexity, LZc) at 1– and 2-hours post 25mg, and significant decreases in alpha power (Figure 1A and B). Additionally, power increases in the gamma band at 2h and decreases in the theta band at 1 and 2h were observed. No changes in EEG recorded activity were observed under 1mg psilocybin (See Table S5 and Figure S6 for these data).

**Figure 1.**
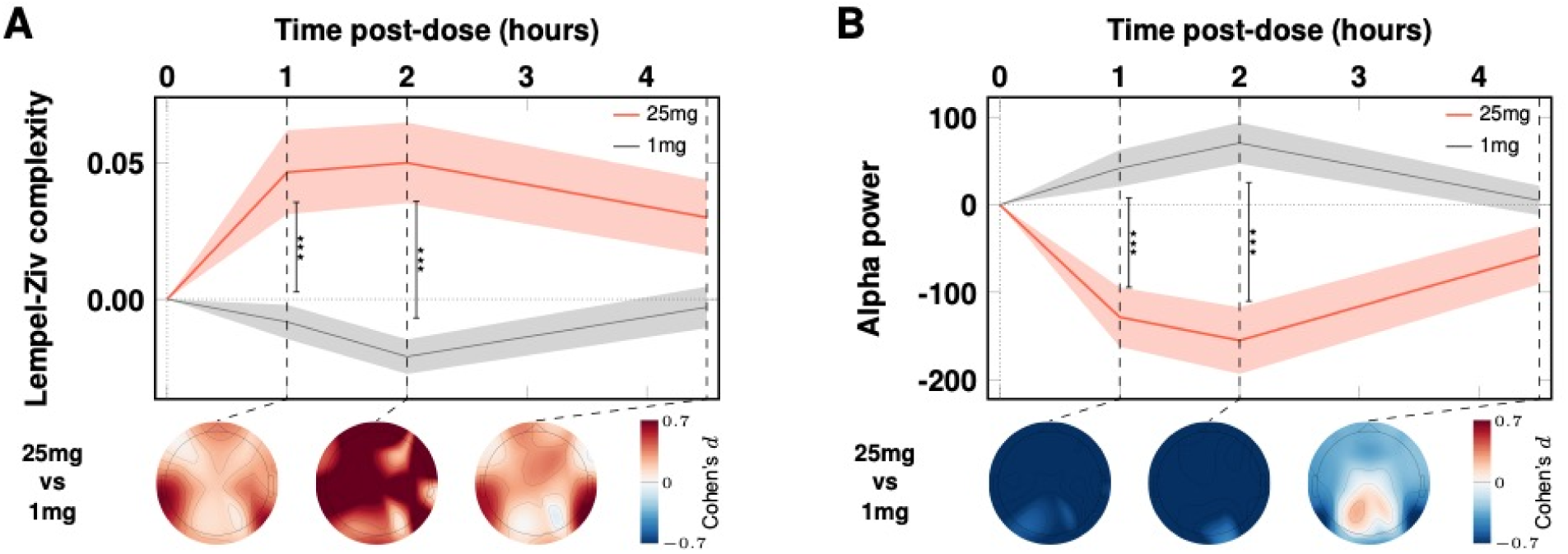
Acute EEG. Significant effects of 25mg (vs 1mg) on brain entropy (LZc) and alpha power. Increases in LZc (A) and decreases in alpha power (B) were maximal at the two-hour timepoint, coinciding with maximum subjective intensity. The red (25mg) and gray (1mg, placebo) timelines correspond to means and standard errors (shaded areas) at 1, 2, and 4.5h post-dose. Vertical bars index contrasts, *** = *p* < 0.001. See Figure S7 for single-subject data.

### Long-term brain changes

#### Anatomical changes (MRI)

Diffusion tensor imaging revealed a significant interaction between ‘*Timepoint*’ (i.e., one factor, three-levels, 1) baseline, 2) 1-month post-1mg, 3) 1-month post-25mg) and mean axial diffusivity (AD) – the diffusion coefficient along the principal diffusion direction, see Figure 2A. ANOVA revealed significant changes in AD for two tracts after 25mg psilocybin, a bilateral prefrontal cortex – striatum tract (PFC-STR; F(2, 48) = 10.30, *p =* 0.005) and a PFC – thalamus tract (PFC-THA; F(2, 48) = 9.51, *p =* 0.008). See Figure S18 for all relevant tracts. Post-hoc t-tests found significant decreases in AD one-month post-25mg vs. one-month post-1mg for the PFC-STR (t(24) = –3.72, *p =* 0.006) and PFC-THA tracts (t(24) = –3.85, *p =* 0.005). Separating by hemispheres for post-hoc t-tests, decreases were apparent in both hemispheres; however, with Bonferroni correction, only decreases in the left hemisphere remained significant (see Figure S19). No significant differences in AD were observed following the 1mg control. With free-water correction (see Figure S21 and section 5.5. of the supplement), consistent AD effects were seen post-25mg psilocybin, and fractional anisotropy (FA) changes also became statistically significant (Figure S21). A tract-based spatial statistics (TBSS) method was also done on request (see Methods) but yielded no significant results.

**Figure 2.**
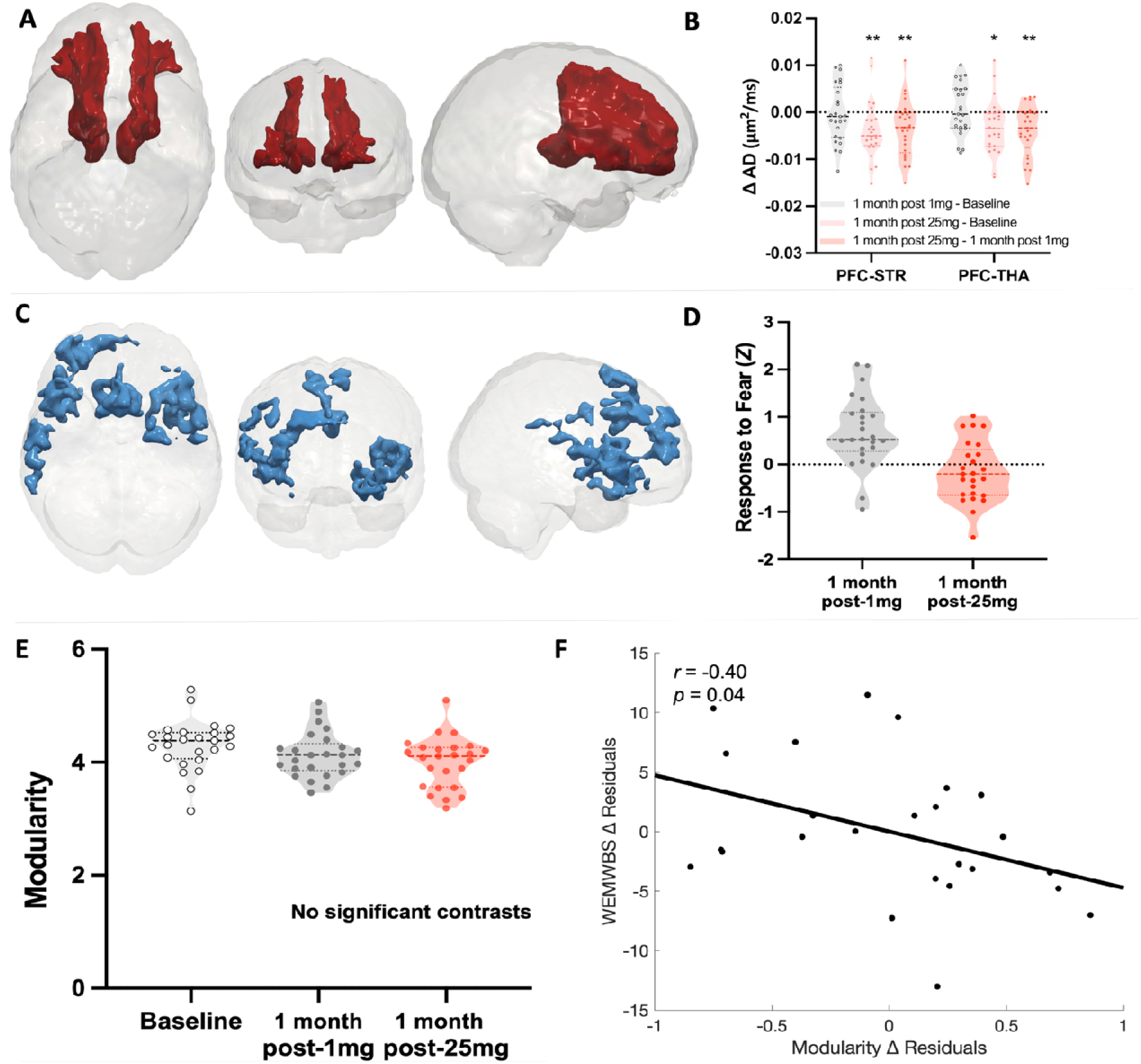
One-month post-25mg psilocybin anatomical and functional brain changes. **A**) **DTI**. Combined bilateral prefrontal-subcortical white-matter tracts where significant decreases in axial diffusivity (AD) were observed one-month post-25mg, Bonferroni corrected. **B**) Violin plot shows decreases in AD with dose; * = *p* < 0.05, ** = *p* < 0.01. See supplement for consistent results with free-water correction (Figure S21) as well as single subject data (Figure S20). **C**) BOLD responses to fearful faces (vs fixation cross) for the key contrast one-month post-25mg vs one-month post-1mg. See Figures S8-9 for baseline and single-subject values. Blue depicts locations of reduced response to fearful faces one-month post-25mg (Z = 2.3, *p <* 0.05, cluster-corrected). **D**) Plots show Z values from the significant clusters shown in C. Note, the delta values for this specific contrast contributes to the correlation matrix shown in Figure 4B; i.e., the variable is labeled ‘Emo-fMRI’ in 4B. **E**) *Brain modularity* at the three timepoints, NS = no contrast was significantly different. **F**) Controlling for *Well-being* prior to 25mg, there was a significant negative correlation between *Modularity Change* and *Well-being Change* in the key contrast of one-month post-25mg vs one-month post-1mg (*r* = –0.4, *p =* 0.04, 2-tailed), i.e., decreased modularity correlated with improved well-being (see also Figure 4B). Note that plotting the residuals in 2F does not highlight the robust improvements in well-being that were seen (note, these changes are evident in 3C). For further DTI and fMRI results, including single subject data, see the supplementary file. Modularity and AD (PFC-STR and PFC-THA tracts merged) change values for the salient one-month post-25mg vs one-month post-1mg contrast also contribute to Figure 4B.

#### Brain response to emotional faces

Whole-brain analyses of the Blood Oxygen Level Dependent (BOLD) fMRI response to emotional face stimuli (happy, fearful, neutral) yielded a large effect (95% *CI* [0.33, 1.34], *d*=0.9) for the salient contrast of one-month post-25mg vs one-month post-1mg psilocybin for fearful faces vs fixation cross, Figures 2C & 2D (see Section 4 of the supplementary file, Figures S8 & S9, for results for happy and neutral faces – and for the post-25mg vs pre-dose baseline contrast; 95% *CI* [0.04, 0.98], *d*=0.3). However, an inclusive (i.e., all factors, all levels) whole-brain analysis, failed to find a significant interaction between emotional face type and *Timepoint* on BOLD response. Explorative amygdala region of interest (ROI) results can be found in the supplement (Figures S10 & S11), along with generalized psychophysiological interaction (gPPI) functional connectivity (FC) findings. Amygdala effects were observed for the left hemisphere and gPPI effects were observed that survived FDR correction for exploring both hemispheres (See Table S12 & Figure S13).

#### Resting-state FC (RSFC)

Explorative ‘resting-state’ (RS) functional connectivity (RSFC) analyses were applied to data from a single 8-minutes eyes-closed, task-free, RS fMRI run. Based on prior hypotheses, four regions were initially chosen for seed-based RSFC analyses: (i) bilateral parahippocampus (PH), (ii) bilateral amygdala, (iii) ventromedial prefrontal cortex (vmPFC), and (iv) subgenual anterior cingulate cortex (sgACC). Two further seeds were examined after peer review: (v) the hippocampus and (vi) the dorsal anterior cingulate cortex (dACC). In ANOVAs across all timepoints, three of the initial four seed did not yield significant results and neither did the further two seeds. However, an increase in amygdala RSFC was evident one-month post-25mg vs one-month post-1mg psilocybin (*M_diff_*=3.68, SE=0.64, *p* < 0.0001; 95% *CI* [2.00, 5.25], *d*=1.2) as well as when post-25mg was contrasted against pre-dose baseline (*M_diff_*=1.49, SE=0.56, *p*=0.046; 95% *CI* [0.02, 2.96], *d*=0.6, Bonferroni-corrected). See Figure S14 for the relevant maps and S15 for single subject data. The robustness of these results was highlighted by an inclusive ANOVA across all timepoints (Figure S16) that found a significant interaction between *Timepoint* and *amygdala RSFC* in consistent regions as those shown in Figure S14 for the key contrast.

#### Network modularity (RSFC)

Previous work highlighted a relationship between improved depressive symptoms and decreased brain network modularity after psilocybin-therapy for depression ^27^. Unlike in this previous work, statistically significant group-average decreases in modularity were not observed here, and neither did we find changes in mass univariate network analyses. More specifically, we found no statistically significant changes in *within* – or *between* network RSFC. However, controlling for *Well-being* before the 25mg session, a correlation was found between *modularity* changes (decreases) one-month post-25mg and contemporaneous changes (improvements) in *Well-being* (*r*= –0.40, *p*=0.04, two-tailed; Figure 2E and F; see also Figure 4B). This relationship is consistent with prior work in depression that found decreased network modularity post-psilocybin correlating with improved depressive symptom severity in two separate trials ^27^.

### Psychological outcomes

#### Acute subjective intensity

An interaction between *Dose* (1mg vs 25mg) and *Time* (0-6h) on the *Intensity* of subjective drug effects was seen (*F*_(8,197)_=57.73, *p <* 0.0001). Greater subjective intensity was reported in relation to 25mg psilocybin compared with both time-zero baseline (all *p* < 0.0001 from 1-5h; *p* < 0.001 at 6h) and the 1mg control (all *p* < 0.0001 from 1-5h; *p =* 0.001 at 6h; Figure 3A). No significant elevation in subjective intensity was observed with 1mg. All, except one, of the participants (94%) rated the 25mg experience as the “*single most unusual state of consciousness”* of their entire life. The remaining person ranked it “*among the top five most unusual experiences*” of their entire life. In contrast, most participants rated their 1mg experience as “*no more unusual than an everyday state of consciousness”* (see Figure S26) reinforcing the impression from the EEG data that the 1mg ‘placebo’ dose was functionally inactive.

**Figure 3.**
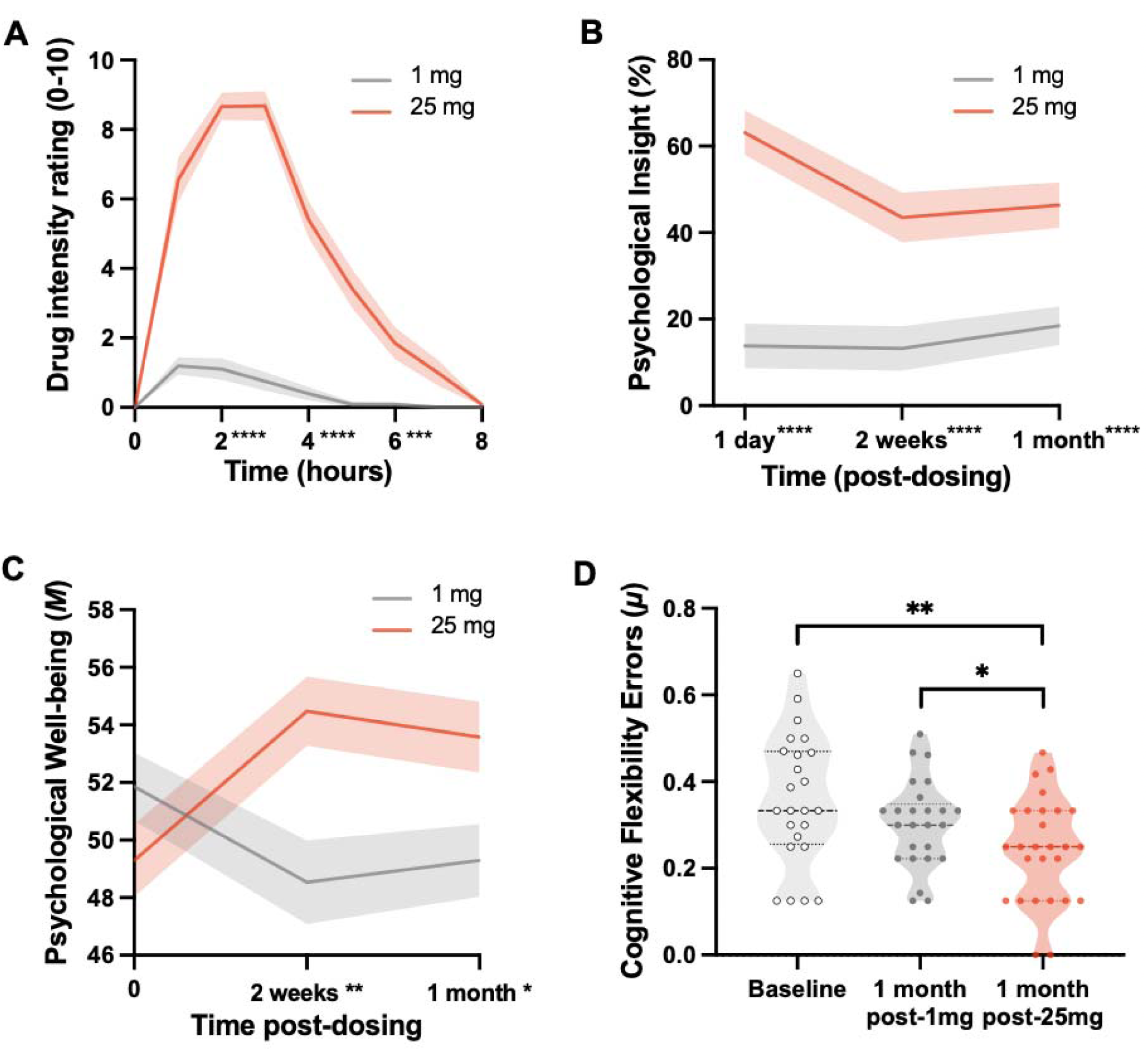
Behavioral analyses. (A) Subjective intensity. A significant interaction between *Dose* (1mg vs 25mg) and *Time* (0-6h) (*F*_(8,197)_ = 57.73, *p <* 0.0001) on *subjective intensity* rated in-session. Post-hoc t-tests, 25mg vs 1mg, *** = *p* < 0.001, **** = *p* < 0.0001. **(B) Psychological insight scale (PIS).** A significant interaction between *Dose* (1mg vs 25mg) and *Timepoint* (one-day, two-weeks & one-month post-dosing) on *psychological insight* (*F*_(2,44)_ = 14.05, *p* < 0.0001); insight scores (PIS) were significantly greater at all timepoints post-25mg vs 1mg (**** = *p* < 0.0001). **(C) Warwick-Edinburgh Mental Well-Being Scale (WEMWBS).** A significant interaction was observed between *Dose* (1mg vs 25mg) and *Timepoint* (pre-dose baseline, two-weeks & one-month post-dose) on *Well-being* (*F*_(2,44)_ = 11.92, *p* < 0.0001); *Well-being* (WEMWBS) increases were significant at two-weeks (*M_diff_* = 5.83, *SE* = 1.54, *p* < 0.01; 95% *CI* [-10.64, –1.02]*, d =* 0.8) and one-month (*M_diff_* = 4.70, *SE* = 1.47, *p* < 0.05; 95% *CI* [-9.29, –0.10]*, d =* 0.6) post-25 mg psilocybin. Post-hoc *t*-tests, 25mg vs 1mg, * = *p* < 0.05, ** = *p* < 0.01. **(D) Intra– and extra-dimensional attentional set-shifting (IDED) & Extradimensional Shift Errors (EDS, Cognitive Flexibility).** Analyses revealed a significant interaction between *Task Performance* and *Timepoint* (F_(7.29,167.67)_ = 2.47, *p* = 0.018; see S23A). Post-hoc tests showed significantly fewer EDS errors in the key contrast of one-month post-25mg vs one-month post-1mg (*M_diff_* = 0.06, *SE* = 0.02, *p* = 0.016*; 95% *CI* [0.00, 0.12], *d* = 0.6), outlier corrected, see Figure S23C for details. Fewer EDS errors reflects faster (correct) detection of rule shifts. Data are expressed as mean ±SEM. T-tests, * = *p* < 0.05, ** = *p* < 0.01. No significant effects of 1mg were observed on any of the statistical tests.

### Long-term psychological changes

#### Psychological insight

Psychological insight was assessed via the ‘psychological insight scale’ (PIS) ^31^. Analyses revealed a robust interaction between *Dose* (1mg vs 25mg) and sub-acute *Timepoints* on *Insight* (*F*_(2,44)_ = 14.05, *p* < 0.0001; Figure 3B). Insight scores were significantly greater after 25mg versus 1mg psilocybin at all sub-acute timepoints— i.e. one-day (*M_diff_* = 50.90, *SE* = 5.51, *p* < 0.0001; 95% *CI* [39.50, 62.29], *d* = 1.9), two-weeks (*M_diff_* = 29.83, *SE* = 4.72, *p* < 0.0001; 95% *CI* [20.08, 39.57], *d* = 1.3) and one-month post-dosing (*M_diff_* = 27.86, *SE* = 4.37, *p* < 0.0001; 95% *CI* [18.90, 36.82], *d* = 1.2), Bonferroni-corrected.

#### Psychological well-being

Psychological well-being was assessed using the Warwick-Edinburgh Mental Wellbeing Scale (WEMWBS), a single dimension, self-report measure of well-being rated with reference to a two-week period ^32^. Analyses revealed a significant interaction between *Dose* (1mg vs 25mg) and sub-acute *Timepoints* on *Well-being* (*F*_(2,44)_ = 11.92, *p* < 0.0001). Bonferroni-corrected *post-hoc* comparisons revealed significantly greater increases in well-being at two-weeks (*M_diff_* = 5.83, *SE* = 1.54, *p* < 0.01; 95% *CI* [-10.64, –1.02]*, d =* 0.8) and one-month (*M_diff_* = 4.70, *SE* = 1.47, *p* < 0.05; 95% *CI* [-9.29, –0.10]*, d =* 0.6) post-25mg psilocybin (vs 1mg), Figure 3C. There were no significant changes post-1mg dose vs (pre any intervention) baseline.

#### Cognitive flexibility

Cognitive flexibility was measured via an intra-dimensional/extra-dimensional (IDED) task, optimised for internet-based delivery ^33,34^. Extradimensional shift (EDS) is an index of participants’ ability to correctly identify rule shifts, and thus, *Cognitive Flexibility*. Participants completed the task at baseline and one-month post-dosing. Greenhouse-Geisser correction was applied due to a violation of sphericity for the interaction term (Epsilon ε = 0.456). Analyses revealed a significant interaction between *Task Discrimination Stage* and *Timepoint* on *Cognitive Flexibility* (*F*_(7.29,167.67)_ = 2.47, *p* = 0.018). FDR-correction of post-hoc comparisons (i.e., for the 9 IDED task phases, Figure 22) plus Bonferroni correction within the salient EDS phase, showed a significant decrease in EDS errors one-month post-25mg vs one-month post-1mg (Figure 3D), indicative of greater cognitive flexibility (*M_diff_* = 0.06, *SE* = 0.02, *p = 0.016*; 95% *CI* [0.00, 0.12], *d*=0.5). Contextualized by no global difference in performance or at IDS, IDR or EDR stages, the EDS change pertains to flexible attentional set as opposed to perseveration or more general problem with processing discriminations / maintaining rules. No IDED changes were observed after 1mg psilocybin. See S22-25 of the supplement for more detailed IDED results.

#### Correlations between long-term change outcomes

The relationship between various contemporaneous (i.e., synchronous in time) changes in brain and behavioral variables observed in the key contrast of one-month post-1mg vs one-month post-25mg was assessed via a correlation matrix that included the relevant contrast values for: 1) *EDS errors* (Cog-flex); 2) *Amygdala-RSFC* (amg-RSFC); 3) *Well-being*; 4) *BOLD responses to emotional faces* (Emo-fMRI); 5) *Brain Network Modularity*; 6) *DTI-measured axial diffusivity* in the merged PFC-THA and PFC-STR tracts (DTI), and 7) *Psychological Insight* (one-month post 1mg vs one-month post 25mg); Figure 4B. Values represent Pearson correlation coefficients, calculated via pairwise linear regressions. Significance was assessed via non-parametric cluster statistics, appropriate for non-independent tests. A significant cluster was found (*p* = 0.006, cluster corrected, coloured cells), implying a nonspurious correlational structure. All of the significant relationships are directionally consistent with the direction of the changes in each outcome for the key contrast of one-month post-1mg vs one-month post-25mg psilocybin.

**Figure 4.**
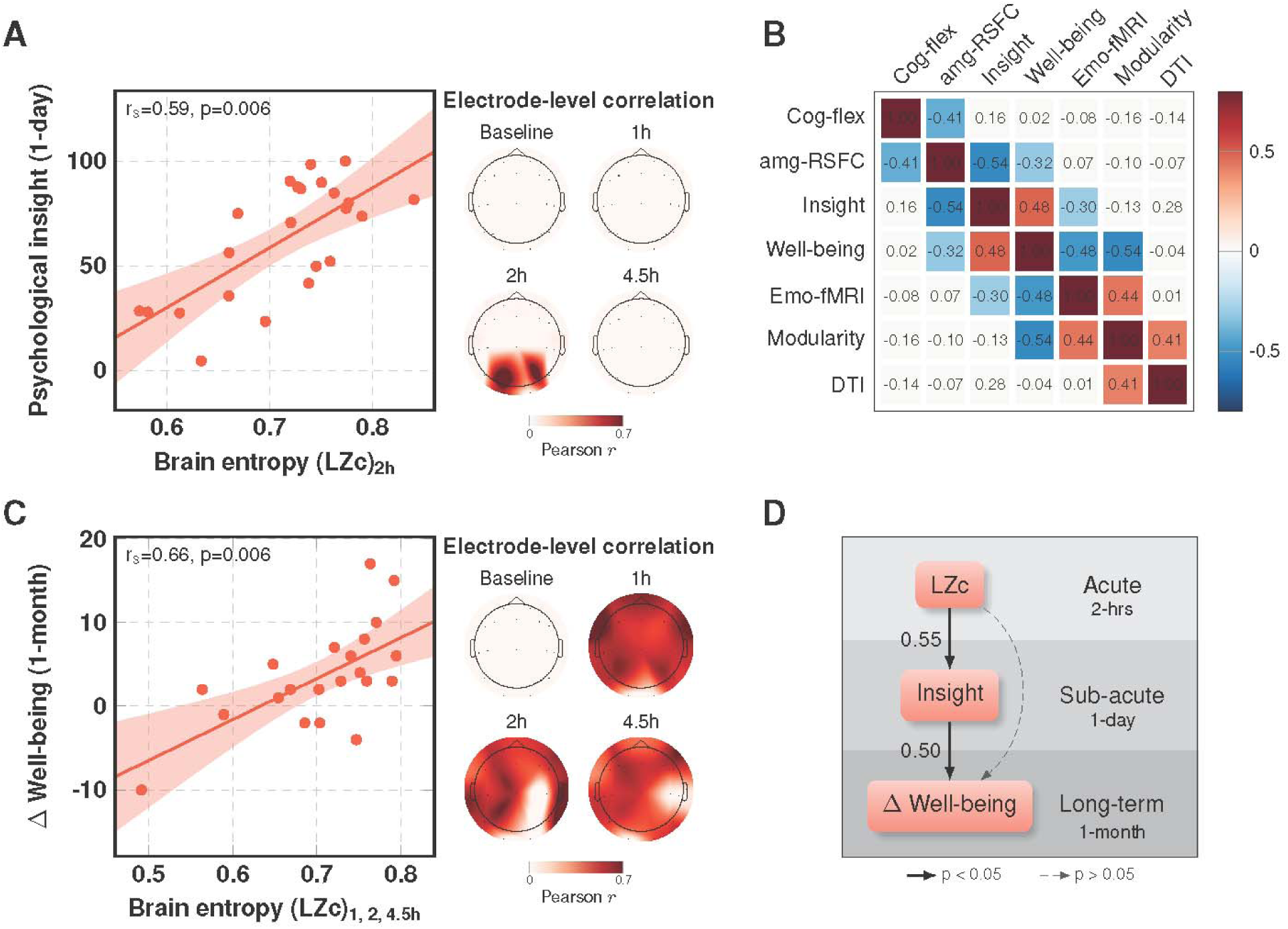
Correlational and predictive modelling. **A**) A cross-validated predictive model revealed a significant positive correlation between acute Lempel-Ziv complexity (LZc, or ‘brain entropy’) for P3, P4, and O1 electrodes, and psychological insight, measured one-day post-25mg. Inset shows electrode-level relationship between LZc (brain entropy) at 2h post-dose (when subjective effects were intense) and next-day insight (see also 3.4 in the supplement for a validation of this result using data from all sensors). **B)** Correlation matrix examining relationships between functional and anatomical MRI metric values and contemporaneous psychological outcomes. Coloured cells represent significant Pearson correlations, corrected for multiple comparisons (non-parametric cluster test, p = 0.006). Red/pink cells reflect significant positive correlations, blue cells reflect significant negative correlations. All relationships are directionally consistent with the original direction of changes. All values derive from the key one-month post-1mg versus one-month post-25mg contrast. **C)** Model predicting one-month well-being increases directly from acute increases in brain entropy at all three post-dosing timepoints. Plot shows electrode-level relationship with (average) data from the 3 post-dose timepoints, i.e., 1-, 2-, and 4.5-hours post-dose (see also 3.4 in the supplement for a validation of the result using data from all sensors). **D**) Model shows that increased well-being at one-month post-25mg can be explained by increased brain entropy (LZc) across all sensors, but also that the strongest path entails brain entropy first predicting next-day *Insight*, which then, in-turn, predicts improvements in *Well-being* (1-month). The values between the arrows in 4D represent normalized regression coefficients. Relationships were statistically significant at p < 0.05.

#### Acute brain entropy predicts increased psychological insight and improved well-being

In two data driven analyses, the informational entropy of spontaneous cortical activity (*LZc*, EEG) under 25mg psilocybin predicted both next-day psychological insight (Figure 4A) and improvements in psychological well-being at one-month post-25mg psilocybin (Figure 4C). In the former case, LZc increases within posterior electrodes (P3, P4, and O1) recorded during 2-hours post-25mg, predicted psychological insight scored one-day later (r_s_ = 0.59, p = 0.006, Figure 4A). In the latter case, LZc predicted improved psychological well-being scores (one-month post-25mg versus one-month post-1mg) directly from most sensors at all three post-dose timepoints (i.e., at 1, 2, and 4.5 hours; *r_s_* = 0.66, *p* = 0.006, Figure 4C). No such predictive relationships were found for the 1mg condition. Note that regressions were done in a data-driven way to identify— via cross-validation — the most sensitive sensors for the relationship i.e., those with the greatest predictive capacity. Repeating these analyses using all sensors contributing equally produced consistent results (see 3.4. in the supplement).

A final mediation analysis (Figure 4D) was performed using linear models that included key variables at three timepoints: (i) whole-brain LZc 2h post-25mg psilocybin (i.e., averaged signal across all sensors at this timepoint), (ii) *Insight* measured one-day post-25mg, and (iii) *Well-being* changes one-month post-25mg. This analysis revealed that acute *LZc* was a significant predictor of next-day *Insight* (β = 197, SE = 67, p = 0.008, normalised β = 0.55). Both *LZc* (β = 41.8, SE = 14.2, p = 0.008, normalised β = 0.54) and *Insight* (β = 0.11, SE = 0.04, p = 0.01, normalised β = 0.50) were significant predictors of increased *Well-being* at one-month; however, the effect of *LZc* on *Well-being* became statistically insignificant if *Insight* was added to the model (β = 29.6, SE = 17.2, p = 0.10, normalised β = 0.38). These results satisfy the criteria for statistical mediation by ^35^ – as well as more recent recommendations ^36^ – implying that the relationship between acute LZc and improved well-being at one-month post-25mg was significantly mediated by sub-acute psychological insight.

## DISCUSSION

The present work sheds important new light on human brain changes after first-time high-dose psilocybin. The high-dose session constituted the first-ever psychedelic experience for all of the study participants. All except one participant rated their high-dose experience with psilocybin as the *single most unusual conscious state* of their entire lives, and the single exception rated it within their *top-five most unusual*. Significant changes in brain function and anatomy were evident from one-hour to one-month after high-dose psilocybin. No acute or long-term changes were apparent after a 1mg ‘placebo’ dose. Empirical modeling highlighted the predictive power of increased brain entropy under high-dose psilocybin, implying its role in manifesting not just the characteristic ‘psychedelic’ effects of psilocybin, but also its longer-term mental-health impact. More specifically, improved well-being could be predicted directly from acute increases in brain entropy as early as one-hour post dosing. Prediction could also be done through a sequence where increased brain entropy first predicted next day psychological insight, which then predicted the one-month improvements in well-being. If non-pharmacological variables are standardized as ‘psychologically supportive’ ^37^, these results imply that human brain changes as early as one-hour into a 25mg psilocybin experience— and that seem closely related to the subjective ‘trip’— can predict mental health improvements one-month later.

Decreased axial diffusivity in prefrontal-subcortical tracts one-month after participants’ first-ever high-dose of a psychedelic could be construed as potential anatomical ‘neuroplasticity’, echoing earlier in-vivo animal studies following single-dose psilocybin that reported increases in synaptic spine formation in female mice^5^ and synaptic density in pigs^6^. However, interpretation of the axial diffusivity changes is complex, not least due to crossing fiber-related confounds^38^. The diffusion-weighted signal can change due to neurofibril or glial growth, altered myelination, axon density, membrane permeability or extracellular fluid ^39^. Decreases in axial diffusivity have been observed with meditation^40^, healthy neurodevelopment^41,42^ and learning^43,44^, but also with axonal injury^45,46^, ageing and related pathology^47^.

5-HT2AR agonism has been associated with *in vivo* dendro-architectural changes in adult mouse brain ^5^, axonal development in embryonic mouse brain ^48^, and oligodendrocyte changes in rodent brain tissue *in vitro* ^49^. The high density of 5-HT2ARs in the human prefrontal cortex ^50^ and greater number of cortical compared with subcortical terminals^51^ suggest a PFC 5-HT2A receptor locus of action for the observed diffusivity changes. The DTI results were robust to free-water correction, and the AD decreases correlated with decreased modularity at one-month post-25mg psilocybin, lending support to the inference that functionally relevant microstructural changes had occurred. Further research using multi-shell sequences is needed to disambiguate the current findings and inform on their robustness and replicability. Until then, serious caution is advised when interpreting the biological basis of these DTI findings. If the effects *are* microstructural, however, then decreases in AD and FA exclusively after the 25mg dose might tentatively relate to two types of change: 1) the pruning of weak or redundant connections; 2) neurogenesis with under-myelinated axons.

Decreased EEG alpha power, a well-replicated effect of psychedelics, is linked with cortical disinhibition ^52^ and increased brain entropy under psychedelics appears to be dose-dependent^53^ and has been associated with acute and sub-acute psychological changes ^54,55^. Decreased cortical alpha power and increased signal entropy are robust and reliable markers of the acute action of psychedelics in humans ^20,21,55–57^. Future research in larger samples could test whether acute EEG changes, induced by 5-HT2AR agonist psychedelics, can predict downstream anatomical neuroplasticity ^3^ or therapeutically-relevant functional brain changes ^27^. However, we failed to find a relationship between acute increases in entropic brain activity and the downstream DTI changes.

Compared with other outcomes in this study, the fMRI-measured brain changes assessed one-month post 25mg were relatively modest; for example, explorative ROI and network-based RSFC analyses (see supplement and methods), including network modularity, yielded largely non-significant results, albeit with some exceptions, including brain responsivity to facial expressions, amygdala RSFC, and a statistically significant *modularity change* to *well-being improvement* correlation (Figure 2C,D,F and Figures S8-17 in the supplement) as well as *modularity change* to *axial diffusivity change* (Figure 4B). Previous work assessing post psilocybin therapy changes in brain function in individuals with depression have revealed more robust changes ^16,27–29^. For example, in two separate trials of psilocybin for depression, decreased brain network modularity was seen that correlated with or predicted improvements in symptom severity post-treatment ^27^ (see also ^58^). Doss et al. found similar effects with psilocybin-therapy for depression that they labelled increased “neural flexibility” ^16^. In the present study, changes in network modularity failed to reach statistical significance (Figure 2E) but we did see a *network modularity change* vs *mental health change* relationship (Figure 2F, 4B) that was directionally consistent with previous work, i.e., decreased modularity correlated with improved mental health ^27^.

Reductions in amygdala and salience-network responses to fearful emotional stimuli post-25mg psilocybin were observed in the present study (Figure 2C&D and S8-11) and these are broadly consistent with prior work in healthy volunteers ^26^. Psychophysiological interaction results were observed for the bilateral amygdala within the face paradigm (see S12-13). Since these analyses were included on request, we refrain from a detailed discussion here but point the reader to a relevant discussion in the supplementary file. Briefly, we note some similarities and differences between a prior study examining amygdala PPI responses to emotional faces post psilocybin-therapy for depression ^59^. Also note, S10&11 (happy > neutral and fear) and two prior psilocybin studies in depression ^28,60^ for evidence of *augmented* brain reactivity to emotional face stimuli post-psilocybin-therapy, most reliably in the non-negative emotional domains – i.e., happy and neutral faces. These latter results could be interpreted as consistent with the re-opening critical periods for social learning ^61^. However, some have questioned the reliability of brain responsiveness to emotional faces as a biomarker of therapeutic action ^62^.

The present study’s fMRI results suggest that enduring functional brain changes post psilocybin are less robust and reliable in healthy versus mentally unwell populations e.g., ^27^. If this principle is reliable, it could imply that greater baseline atypicality in the clinical populations, primes for a more robust remediating change via psilocybin. However, given the reliability of enduring psychological changes post psilocybin in healthy samples ^18,19^, it remains plausible— if not likely— that functional brain changes *do* occur in healthy populations but their detection is dependent on— or sensitive to— experimental modality, metric and parameter choices ^27,29,63^. In short, we may have not yet found a sufficiently sensitive assay to detect true functional brain changes.

The present multi-modal neuroimaging study in healthy participants sheds new light on the brain effects of first-time high-dose psychedelic use and the therapeutic action of psilocybin-therapy, suggesting, for the first time, how therapeutically relevant effects (i.e., improved well-being) can be forecast via an *acute* human brain action, i.e., entropic brain activity, that is known to relate to the subjective ‘trip’ ^55^. Recent evidence suggests that this characteristic effect of psychedelics is dose-dependent and somewhat exclusive to this category of drug versus psychoactive stimulants and cannabinoids ^53^. Long-term improvements in well-being were predicted by the acute increases in entropic brain activity, temporally coinciding with the psilocybin’s acute subjective effects. Results support a role for psychological insight in mediating the causal association between increased entropic brain activity and potentially enduring improvements in well-being. These and prior results highlight psychological insight ^31^ and entropic brain effects ^64,65^ as important to the action of psychedelic therapy. Possible white-matter changes, as well as improvements in well-being and cognition were also observed in this study. All acute and enduring psychological and neurobiological effects were exclusive to the high-dose psilocybin condition.

Justified by prior evidence of enduring psychological changes post-psychedelics ^9,10^, this study used a fixed-order cross-over design. Pairwise contrasts on all the study’s main outcomes showed no evidence of acute brain or behavioural effects or changes one-month post-1mg; however, order confounds cannot be entirely discounted. Between-subject confirmatory studies are now required to examine the reliability of these novel findings.

## Supporting information

Supplementary results

Methods section

## ACKNOWLEDGEMENTS

Funded by philanthropic donations to the Centre for Psychedelic Research, Imperial College London, founded in April 2019. TL received an MRC-DTP Studentship, HMD received an Imperial College London President’s PhD Scholarships. RCH was supported by the Alex Mosley Charitable Trust between 2015 and April 2019. From April 2019 to July 2021, RCH was supported by donations to the Centre for Psychedelic Research at Imperial College London, upon its creation; and from July 2021 to present, RCH has been supported by The Ralph Metzner Distinguished Professorship at University of California San Francisco. Some further support for this study was provided by The Beckley Foundation. Infrastructure support was provided by the NIHR Imperial Biomedical Research Centre and the NIHR Imperial Clinical Research Facility. The views expressed are those of the author(s) and not necessarily those of the NHS, the NIHR or the Department of Health and Social Care.

We thank Manca Peskar, Victoria Nygart, Hannes Kettner and Mellissa Shukuroglou for their invaluable assistance on study visits; Dr Roberta Murphy and Dr Jonny Martell for clinical oversight; Dr Rosalind Watts for advice, training and support on guiding and integration procedures; Dr Robin Tyacke for controlled drug oversight; Joe Peill, Gregory Cooper, Katie Trinci and Pablo Mallaroni for their support with data analysis; and Bruna Giribaldi for advice and support.

## AUTHOR CONTRIBUTIONS

RCH designed the study and wrote the original and revised manuscripts. RCH and DJN provided oversight. TL coordinated the study, led the study visits, collected the data, created charts and plots, and worked with RCH to compile the paper and revisions. MS coordinated the study to completion, leading study visits and collecting data for the final set of participants. LE, LR, CT and FR supervised dosing sessions and performed data collection. LE and DE provided medical cover and assisted with study visits. TL analyzed the behavioral data and conducted regression and correlation analyses. FR and PMM ran the clustering analyses of long-term changes, completed the mediation analysis, and created the corresponding figures (Figures 4A-D). PMM and FR also created Figure 1A-B. LK analyzed the diffusion MR data. LR, BAP, CT, MG, and MBW analyzed the functional MR data. LK created the diffusion images and RCH organized them into plots. FR, PMM, FS, CT, MS, and HD analyzed the EEG data. AH and WT guided our use of the IDED, which HD analyzed, and TL created charts for. BAP and RJZ led the PPI analyses and created the relevant charts. All authors were given the opportunity to provide edits of the manuscript and accepted the final draft.

