## Supplementary results for "Human brain changes after first psilocybin use"

**SUPPLEMENTARY MATERIAL**

**Contents**

**1. Background**

1.1. Table S1. Reference table for (to our knowledge) all previous psychedelic studies involving sub or post-acute neuroimaging or behavioural outcomes.

**2. Study design and demographics.**

2.1. Recruitment, screening, and eligibility criteria.

2.2. Figure S2. Study timeline of interventions.

2.3. Table S3. Schedule of interventions.

2.4. Table S4. Demographic data.

**3. EEG outcomes**

3.1. Table S5. EEG modeling of complexity and power changes in frequency bins and interaction between Dose and Timepoint.

3.2. Figure S6. Sensor-level EEG maps for LZ and power in frequency-bins.

3.3. Figure S7. Single subject changes in LZ and alpha power.

3.4 Additional controls pertaining the results of the data-driven analysis of LZ.

**4. fMRI outcomes**

4.1. Figure S8. fMRI BOLD responses to emotional faces (all face types) and all time points. Data drawn from 25mg vs 1mg cluster.

4.2. Figure S9. Response to faces with single subject values shown.

4.3. Amygdala (ROI) response to emotional faces.

4.4. Figure S10. Left amygdala BOLD response to emotional faces of various types.

4.5. Figure S11. Right amygdala BOLD responses to emotional faces.

4.6. Table S12. Amygdala psychophysiological interaction analysis: One-month post 1mg vs one-month post-25mg.

4.7 Figure S13. Amygdala psychophysiological interaction analysis: One-month post 1mg vs one-month post-25mg.

4.8. Figure S14. Bilateral amygdala BOLD RSFC analysis on key contrast.

4.9. Figure S15. Single subject data. Bilateral amygdala BOLD RSFC.

4.10. Figure S16. Whole-brain naïve ANOVA across all timepoints.

4.11. Validation analysis: in-scanner anxiety: amygdala RSFC analysis.

4.12. Resting State Networks (RSN) – Within and Between Networks RSFC.

4.13. Figure S17. Modularity with single subject values shown.

**5. DTI outcomes**

5.1. Figure S18. DTI: tracts where changes were observed.

5.2. Figure S19. Axial diffusivity values without free-water correction. Left hemisphere.

5.3. Figure S20. Axial diffusivity values with single subject datapoints shown.

5.4. Figure S21. Fractional Anisotropy (FA) with free-water correction.

5.5. Free-water correction values for axial diffusivity.

**6. Psychological outcomes**

6.1 Detailed description of Intradimensional/extradimensional (IDED) task.

6.2. Figure S22. IDED task phase 1-9.

6.3. Figure S23. IDED task: Cognitive flexibility (more granular) findings.

6.4. Figure S24. EDS errors. Single subject data.

6.5. Table S25. IDED task errors per criterion.

6.6. Figures S26. Adapted version of ‘Persisting Effects Questionnaire’ of Griffiths et al. 2006.

**7. References**

**1. Background**

1.1 **Table S1.**

| **Reference and URL link** | Brief description |
| --- | --- |
| Roseman L, Demetriou L, Wall MB, et al. (2018) Increased amygdala responses to emotional faces after psilocybin for treatment-resistant depression. *Neuropharmacology* 142: 263-269  <https://doi.org/10.1016/j.neuropharm.2017.12.041> | Increased amygdala responses to emotional faces after psilocybin for treatment-resistant depression (this group's work) |
| Mertens LJ, Wall MB, Roseman L, et al. (2020) Therapeutic mechanisms of psilocybin: Changes in amygdala and prefrontal functional connectivity during emotional processing after psilocybin for treatment-resistant depression. *Journal of Psychopharmacology* 34(2): 167-180.  <https://doi.org/10.1177/0269881119895520> | Decreased amygdala functional connectivity 1 day after psilocybin during a task in patients with depression (this group's work) |
| Stroud JB, Freeman TP, Leech R, et al. (2018) Psilocybin with psychological support improves emotional face recognition in treatment-resistant depression. *Psychopharmacology (Berl)* 235(2): 459-466.  <https://doi.org/10.1007/s00213-017-4754-y> | Enhancements of emotional processing 1 day after psilocybin in patients with depression (this group's work) |
| Doss MK, Považan M, Rosenberg MD, et al. (2021) Psilocybin therapy increases cognitive and neural flexibility in patients with major depressive disorder. *Translational Psychiatry* 11(1): 574.  <https://doi.org/10.1038/s41398-021-01706-y> | Enhanced cognitive flexibility 1 week after psilocybin in patients with depression |
| Barrett FS, Doss MK, Sepeda ND, et al. (2020) Emotions and brain function are altered up to one month after a single high dose of psilocybin. *Scientific Reports* 10(1): 2214.  <https://doi.org/10.1038/s41598-020-59282-y> | Decreased amygdala activation 1 week after psilocybin and no changes in within- and between-network connectivity in a healthy population (like the current study) |
| Smigielski L, Scheidegger M, Kometer M, et al. (2019) Psilocybin-assisted mindfulness training modulates self-consciousness and brain default mode network connectivity with lasting effects. *Neuroimage* 196: 207-215.  <https://doi.org/10.1016/j.neuroimage.2019.04.009> | fMRI 2 days after psilocybin |
| McCulloch DEW, Madsen MK, Stenbæk DS, et al. (2021) Lasting effects of a single psilocybin dose on resting-state functional connectivity in healthy individuals. *Journal of Psychopharmacology* 36(1): 74-84.  <https://doi.org/10.1177/02698811211026454> | fMRI 1 week and 3 months after psilocybin |
| Mason NL, Kuypers KPC, Reckweg JT, et al. (2021) Spontaneous and deliberate creative cognition during and after psilocybin exposure. *Translational Psychiatry* 11(1): 209.  <https://doi.org/10.1038/s41398-021-01335-5> | fMRI acutely but also cognitive testing 1 week after psilocybin |
| Mason NL, Mischler E, Uthaug MV, et al. (2019) Sub-Acute Effects of Psilocybin on Empathy, Creative Thinking, and Subjective Well-Being. *Journal of Psychoactive Drugs* 51(2): 123-134.  <https://doi.org/10.1080/02791072.2019.1580804> | Cognitive testing 1 day and 1 week after psilocybin |
| Smigielski L, Kometer M, Scheidegger M, et al. (2019) Characterization and prediction of acute and sustained response to psychedelic psilocybin in a mindfulness group retreat. *Scientific Reports* 9(1): 14914.  <https://doi.org/10.1038/s41598-019-50612-3> | Predicting the sustained response of psilocybin |
| Sampedro F, de la Fuente Revenga M, Valle M, et al. (2017) Assessing the Psychedelic “After-Glow” in Ayahuasca Users: Post-Acute Neurometabolic and Functional Connectivity Changes Are Associated with Enhanced Mindfulness Capacities. *International Journal of Neuropsychopharmacology* 20(9): 698-711.  <https://doi.org/10.1093/ijnp/pyx036> | fMRI 1 day after ayahuasca |
| Pasquini L, Palhano-Fontes F and Araujo DB (2020) Subacute effects of the psychedelic ayahuasca on the salience and default mode networks. *Journal of Psychopharmacology* 34(6): 623-635.  <https://doi.org/10.1177/0269881120909409> | fMRI1 day after ayahuasca |
| Uthaug MV, van Oorsouw K, Kuypers KPC, et al. (2018) Sub-acute and long-term effects of ayahuasca on affect and cognitive thinking style and their association with ego dissolution. *Psychopharmacology (Berl)* 235(10): 2979-2989.  <https://doi.org/10.1007/s00213-018-4988-3> | Cognitive testing 1 day and 4 weeks after ayahuasca |
| Kiraga MK, Mason NL, Uthaug MV, et al. (2021) Persisting Effects of Ayahuasca on Empathy, Creative Thinking, Decentering, Personality, and Well-Being. *Frontiers in Pharmacology* 12:721537.  <https://doi.org/10.3389/fphar.2021.721537> | Cognitive testing 1 day and 1 week after ayahuasca |
| Bouso JC, Palhano-Fontes F, Rodriguez-Fornells A, et al. (2015) Long-term use of psychedelic drugs is associated with differences in brain structure and personality in humans. *European Neuropsychopharmacology* 25(4): 483-492.  <https://doi.org/10.1016/j.euroneuro.2015.01.008> | Another study, albeit observational in nature, that reported structural (grey matter) changes from a psychedelic i.e., ayahuasca use |
| Murphy-Beiner A and Soar K (2020) Ayahuasca’s ‘afterglow’: improved mindfulness and cognitive flexibility in ayahuasca drinkers. *Psychopharmacology (Berl)* 237(4): 1161-1169.  <https://doi.org/10.1007/s00213-019-05445-3> | Enhanced cognitive flexibility 1 day after ayahuasca in an ayahuasca using population |
| Halpern JH, Sherwood AR, Hudson JI, et al. (2005) Psychological and cognitive effects of long-term peyote use among Native Americans. *Biological Psychiatry* 58(8): 624-631.  <https://doi.org/10.1016/j.biopsych.2005.06.038> | No effect on cognitive flexibility approximately 1 month after mescaline in a mescaline-using population |
| Wießner I, Olivieri R, Falchi M, et al. (2022) LSD, afterglow and hangover: Increased episodic memory and verbal fluency, decreased cognitive flexibility. *European Neuropsychopharmacology* 58: 7-19.  <https://doi.org/10.1016/j.euroneuro.2022.01.114> | Impaired cognitive flexibility 1 day after LSD |

2. **Study design and demographics**
   2.1. **Recruitment, screening, and eligibility criteria**

Participants (*N*=28) were recruited via an online advertisement on the Centre for Psychedelic Research website. All participants initiated contact to gain their place on the study. After an initial telephone screen, participants were formally screened in the National Institute for Health Research/Wellcome Trust Imperial Clinical Research Facility (ICRF) to determine eligibility. Participants were emailed the participant information sheet outlining the nature, purpose and risks of the study in sufficient time for them to read it prior to the screening visit. Participants were encouraged to ask questions about the information provided before giving their informed consent.

During screening, participants underwent physical and mental health assessments. Physical assessments included: (i) an alcohol breathalyser test, (ii) a urine screen for drugs of abuse and pregnancy (where applicable), (iii) an electrocardiogram (ECG), (iv) routine blood tests, and (v) blood pressure, heart rate, height and weight recordings. Mental health assessments included (i) a neurological examination and (ii) a standard psychiatric interview, the Mini-International Neuropsychiatric Interview (MINI) version 5. Participants provided their full medical history and declared previous drug use. Screenings typically lasted 2.5 hours. After the retrieval of screening results, eligible participants were enrolled in the study and invited to return for subsequent visits.

*Inclusion criteria*

- Physically and mentally healthy
- Males & females
- 18-85 years old
- Good command of the English language
- No prior experience with psychedelic drugs

*Exclusion criteria*

- Current or previously diagnosed psychiatric disorder
- One or more immediate family members with a current or previously diagnosed psychotic disorder
- Medically significant condition that renders them unsuitable for the study (e.g., diabetes, severe cardiovascular disease, hepatic or renal failure etc.)
- Show MR contraindications (e.g., metal implants, pacemakers, claustrophobia etc.)
- Prior experience with a psychedelic drug
- Blood or needle phobia
- Show a positive pregnancy test at screening or during the study
- Excessive use of alcohol or other drugs (determined by study medic)
- No email access
- Inadequate grasp of English language
- Currently or very recently (within 6 months) involved in other studies
- Use of alcohol within 24 hours or other drugs within 7 days of scans and doses

**2.2. Figure S2. Study timeline of interventions**

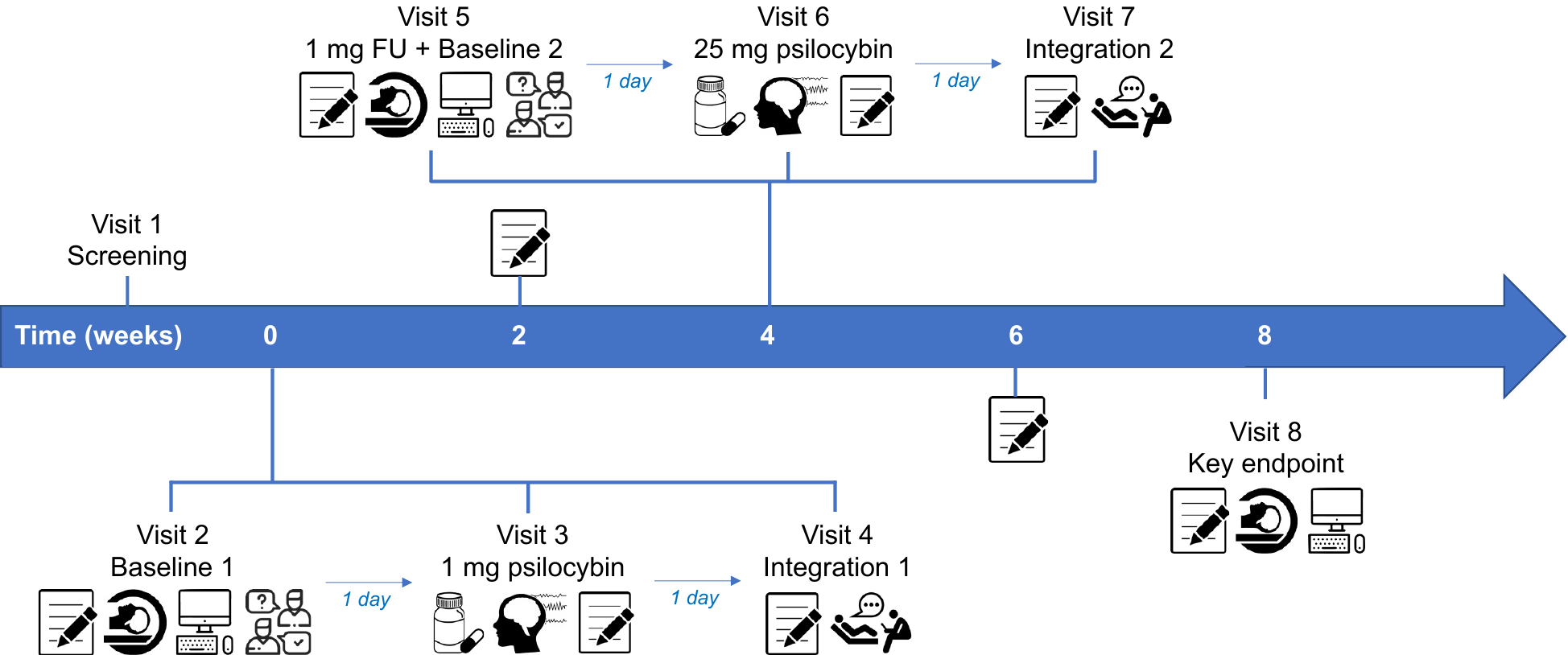

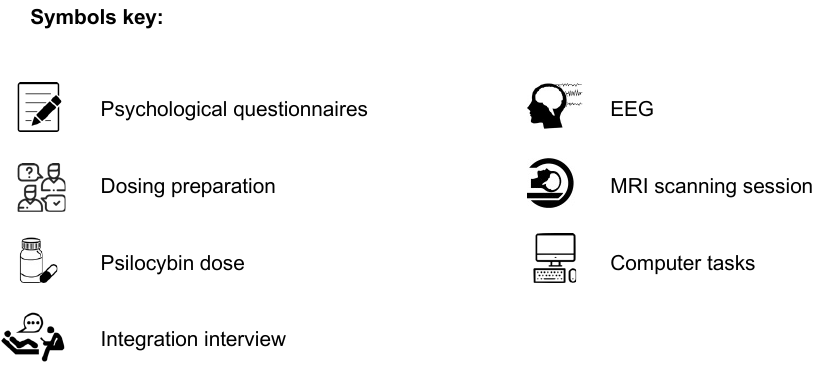

**Figure S2**. This figure shows the timeline of events in this study. Visit 2 was the true pre-intervention baseline; 1mg psilocybin dosing occurred one day later on visit 3. One-month elapsed (i.e., typically 4 weeks or just over) until visit 5, which was one day prior to 25mg psilocybin dosing (visit 6); visit 5 acted as a follow-up for the control dose of 1mg as well as a second pre-intervention (25mg psilocybin) baseline for assessing the effects of 25mg psilocybin. A further month elapsed before the key endpoint, visit 8, one-month post-25mg psilocybin. Psychological assessments were conducted on all visits as well as remotely 2 weeks after each dose. EEG recordings were completed acutely during dosing days on visits 3 and 6. Participants returned one day after each dosing day on visits 4 and 7 for a psychological integration session, involving a post-dosing check-up, open listening, and an interview about their experience. MRI scanning and a separate computer task assessing cognitive flexibility was conducted on visits 2, 5 and 8. In brief, visit 2 is baseline 1, visit 5 is baseline 2 and also the follow-up check on potential effects elicited by the control (1mg), and visit 8 is the final follow-up to assess the effects of 25 mg psilocybin.

**2.3. Table S3. Schedule of interventions.**

|  | **PRE-DOSING** | **ACUTE** | **POST-DOSING** | | |
| --- | --- | --- | --- | --- | --- |
|  |  |  | ***1 day*** | ***2 weeks*** | ***1 month*** |
| **Intensity** | X | X |  |  |  |
| **EEG** | X | X |  |  |  |
| **PIS** |  |  | X | X | X |
| **WEMWBS** | X |  |  | X | X |
| **IDED** | X |  |  |  | X |
| **MRI** | X |  |  |  | X |

**Table S3**. Table showing a schedule of interventions. “Pre-dosing” refers to the pre-dosing baseline, which occurred one-day prior to the 1mg dosing session. Acute refers to measures done on the dosing day itself. Post-dosing timepoints are labeled in the Table header. Intensity refers to the subjective intensity of the drug effects verbally reported by the participants each hour after dosing, using a 0-10 scale, where ‘10’ = the most intense effects imaginable. Abbreviations: EEG, Electroencephalography; PIS, Psychological Insight Scale; WEMWBS, Warwick-Edinburgh Mental Well-being Scale; IDED, Intra-dimensional/extra-dimensional task; MRI, Magnetic Resonance Imaging.

**2.4. Table S4. Demographic data**

| **Demographic variable** | ***M*** | ***SD (range)*** |
| --- | --- | --- |
| **Age** (in years) | 40.6 | 8.7 (29-59) |
|  | ***n*** | **%** |
| **Gender**  Male  Female | 16  12 | 57%  43% |
| **Ethnicity**  Caucasian  Undisclosed  Black | 24  3  1 | 86%  11%  3% |
| **Nationality**  British  Other | 21  7 | 75%  25% |
| **Education**  Secondary School Level  University Level | 12  16 | 43%  57% |
| **Employment status**  Full time  Part time  Unemployed | 24  3  1 | 86%  11%  3% |

**Table S4**. Age data expressed as mean ± SD with the age range. The chi-square test was used to determine within-sample differences in categorical variables. Data expressed as frequency count (n) and percentage (%) of sample. Participants (N=28) had an average age of 41 years (*SD=*8.7, range: 29–59) and were balanced in terms of gender (*χ^2^*=0.57, *p=*0.450) and educational attainment (*χ^2^*=0.62, *p=*0.430). All participants were naïve to psychedelic drugs and the majority were British (75%; *χ^2^*=7.00, *p<*0.01) and Caucasian (86%; *χ^2^*=14.57, *p<*0.001) in full-time employment (86%; *χ^2^*=14.29, *p<*0.001).

3. **EEG outcomes**

**3.1. Table S5. Psilocybin induces time-dependent changes in LZ and spectral power**

|  | **Doses (1mg vs 25mg) x timepoints (TPs) in hours (h) post-administration** | | | | | | |
| --- | --- | --- | --- | --- | --- | --- | --- |
|  | **Time**  **1 h** | **Time**  **2 h** | **Time**  **4.5 h** | **Dose x**  **Baseline** | **Dose x**  **1h** | **Dose x**  **2 h** | **Dose x 4.5 h** |
| **β (LZ)** | -0.01 | -0.02 | -0.003 | -0.01 | **0.05** | **0.07** | 0.04 |
| **SE** | 0.01 | 0.01 | 0.01 | 0.01 | **0.01** | **0.01** | 0.01 |
| ***p*** | 0.359 | 0.036 | 0.773 | 0.404 | **4x10^-4** | **2x10^-6** | 0.013 |
| **β (delta)** | 35.8 | 25.3 | -13.4 | -25.4 | -19.2 | 72.2 | -28.7 |
| **SE** | 33.5 | 33.5 | 32.6 | 36.1 | 47.3 | 47.3 | 46.4 |
| ***p*** | 0.287 | 0.452 | 0.682 | 0.482 | 0.685 | 0.129 | 0.538 |
| **β (theta)** | 24.7 | 29.2 | -3.5 | -7.1 | -48.4 | -53.8 | -23.2 |
| **SE** | 14.6 | 14.6 | 14.2 | 14.2 | 20.6 | 20.6 | 20.3 |
| ***p*** | 0.093 | 0.047 | 0.808 | 0.622 | 0.020 | 0.010 | 0.254 |
| **β (alpha)** | 37.9 | 67.5 | 4.7 | 4.5 | **-163.5** | **-223.0** | -68.5 |
| **SE** | 27.1 | 27.1 | 26.4 | 29.4 | **38.3** | **28.3** | 37.6 |
| ***p*** | 0.165 | 0.014 | 0.869 | 0.877 | **4x10^-5** | **4x10^-8** | 0.071 |
| **β (beta)** | 6.51 | 6.94 | -6.04 | -4.42 | -15.57 | -7.66 | -4.18 |
| **SE** | 5.14 | 5.14 | 5.01 | 5.33 | 7.26 | 7.26 | 7.13 |
| ***p*** | 0.207 | 0.179 | 0.230 | 0.408 | 0.034 | 0.293 | 0.559 |
| **β (gamma)** | 2.05 | 1.27 | -6.40 | -3.68 | 7.82 | **18.48** | 5.59 |
| **SE** | 3.72 | 3.72 | 3.63 | 3.94 | 5.26 | **5.26** | 5.17 |
| ***p*** | 0.583 | 0.732 | 0.080 | 0.352 | 0.140 | **6x10^-4** | 0.281 |

**Table S5.** Statistics (estimates, standard error, and uncorrected p-values) of the effects of each timepoint, dose (1mg vs 25mg), and their interaction (rightmost 3 columns), as obtained by a linear mixed model. The model used timepoint and dose as independent, categorical values (with base level at baseline and 1mg, respectively), and a random intercept to account for individual differences. Each set of rows corresponds to a model with a different dependent variable, specifically LZ and spectral power in the delta, theta, alpha, beta, and gamma bands (rows along the vertical). Red font highlighted p-values are the ones that survive Bonferroni correction. D=dose (25 or 1 mg), h=hours after administration. There is a significant interaction between dose (1mg vs 25mg) and timepoint at 1hr and 2hrs on LZ and alpha. There is also a significant interaction of dose and timepoint at 2hrs for gamma.

**3.2.** **Figure S6. Sensor-level EEG maps for LZ and power in 5 canonical frequency bins**

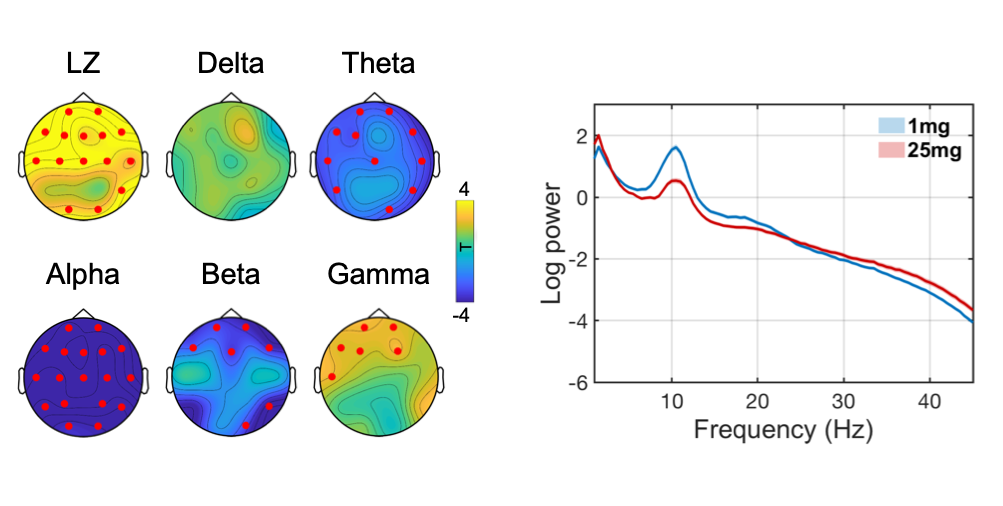

**Figure S6.** Maximum effects of 25mg vs 1 mg of psilocybin in EEG features (LZ and power in frequency bands) occurred 2-hours after administration. Localized scalp changes are marked with red electrodes are shown in the left (p<0.05, two-sided, cluster-corrected). Averaged power spectra of brain activity at 2h, is shown on the right for both 25mg (red) and 1mg (blue) conditions.

**3.3. Figure S7. Single subject changes in LZ and alpha power with 1mg and 25mg psilocybin**

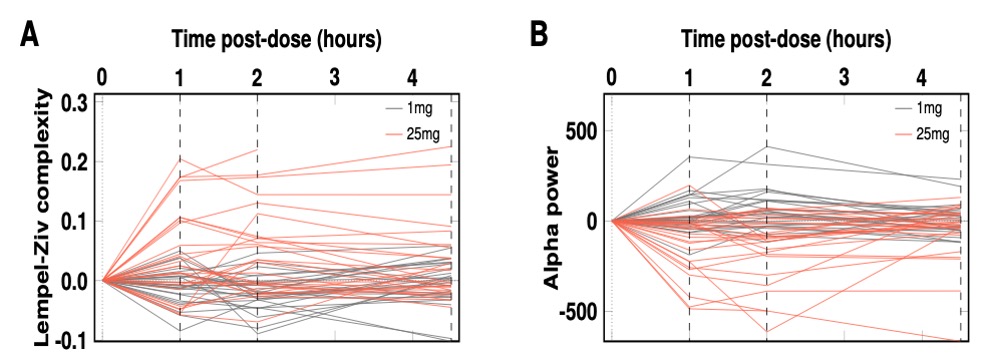

**Figure S7.** Single subject changes in LZ (A) and alpha power (B) from pre-dose (time zero) to the 3 salient post-dose recording timepoints (1hr, 2hrs, 4.5hrs). See Figure 1 of the main manuscript to view the group level data, including variance and relevant statistical tests.

**3.4 Additional controls pertaining the results of the data-driven analysis of LZ.**

Here we report the findings of additional controls related to the analysis of LZ values computed from the EEG data, involving the calculation of the Spearman correlation between simple spatial averages of LZ values across electrodes obtained at different time-points and correcting for multiple comparisons via Bonferroni (see Methods). Results showed that LZ measured at 2hrs post 25mg psilocybin ingestion is significantly correlated with insight reported the day after (r=0.55, p=0.020), while LZ measured at 1hrs (r=0.41, p=0.14) and 4.5hrs (r=0.43, p=0.11) don’t survive correction. These results replicate the sensor-specific results presented in the main paper (Figure 4A) wherein data from the 2hrs timepoint is shown. When running similar analyses between LZ and well-being measured one-month after, results show significant correlations between LZ measured at 1hrs (r=0.58, p=0.010) and 2hrs (r=0.54, p=0.022), but no significant correlations with LZ measured 4.5hrs (r=0.41, p=0.144). These results, obtained by much simpler means, and with a conservative multiple comparison correction, support the validity of the data-driven analyses presented in the main text. All relationships are exclusive to the 25mg dosing session.

4. **fMRI outcomes**

**4.1.	Figure S8. fMRI BOLD responses to emotional faces (all three face types)**

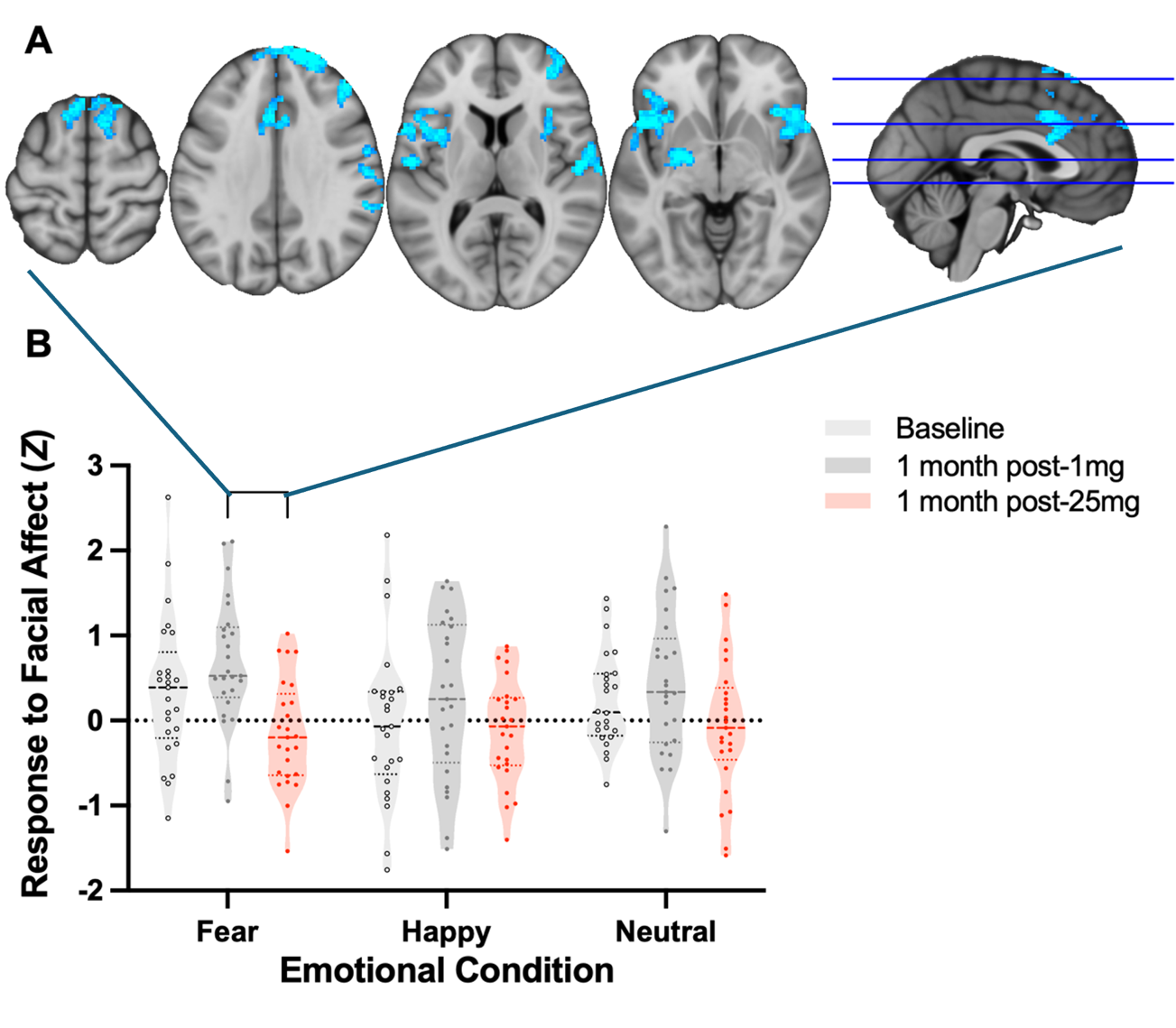

**Figure S8. A**) Whole brain voxelwise analysis revealed significantly reduced BOLD responses to fearful faces one-month after 25mg versus one-month after 1mg (Z = 2.3, p < 0.05). This pattern of reduced responsiveness resembled the so-called ‘Salience Network’. **B**) Violin plots showing BOLD responses to emotional faces of various types derived from the regions shown in Blue in A. Note, as data shown in B is derived from the one-month post-1mg vs one-month post-25mg cluster for Fear vs fixation cross (i.e., the clusters shown in A) - i.e., a test-selective cluster of voxels, we refrain from showing statistical significance tests on the values presented in this figure. Note. An inclusive ANOVA with time and emotional face type as factors failed to yield a significant result and therefore no map for this test is shown.

**4.2.	Figure S9. Response to faces with single subject values shown.**

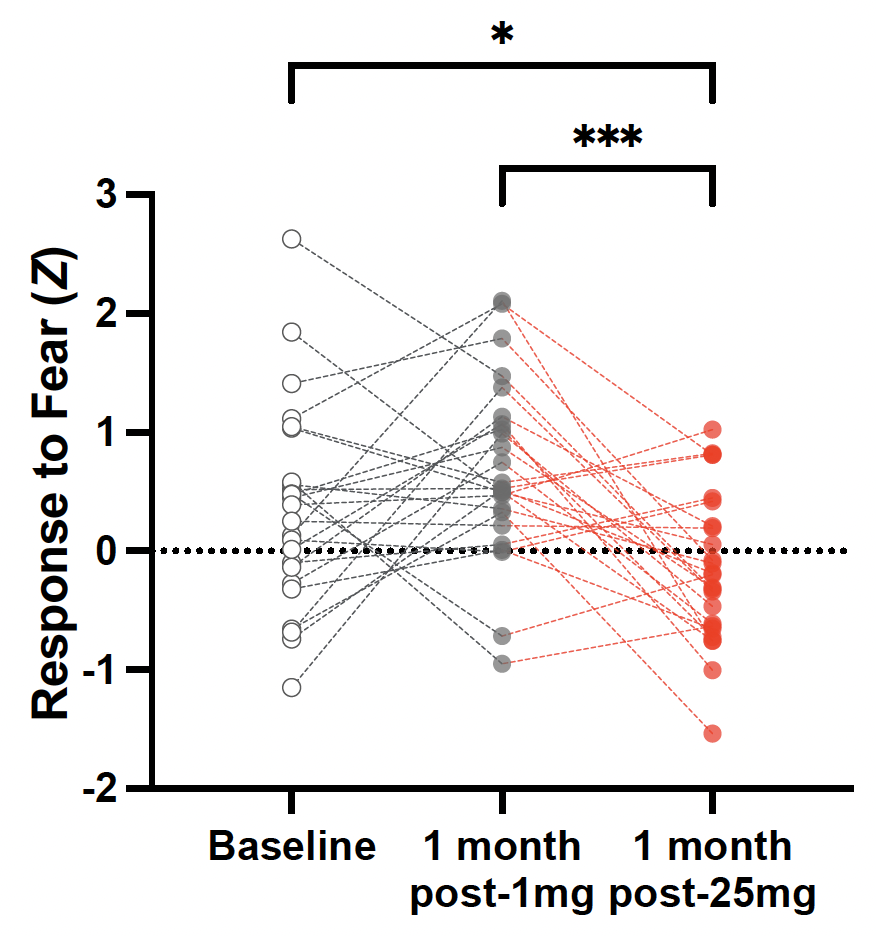

**Figure S9.** Response to fear with single subject datapoints shown. These values are drawn from the cluster that was significant in the 25mg vs 1mg contrast (i.e., the cluster shown in Figure S7A - also the left-most plot of S7B). We refrain from making statistical tests on these data.

**4.3.	Amygdala (ROI) response to emotional faces**

ROI analyses of the amygdala response (amygdala mask based on Harvard-Oxford atlas, with threshold > 50) to facial stimuli yielded a statistically significant interaction between emotional condition and time in the left amygdala (*F*_(8,192)_=3.32, *p*=0.001), particularly within a distinct cluster (*F*_(8,192)_=3.95, *p*<0.001); this cluster within the left amygdala is shown in Figure S7A. FDR corrected post-hoc comparisons across all timepoints and conditions showed a significant decrease in BOLD response to fearful facial stimuli one-month after 25 mg psilocybin (*M_diff_*=-0.81, SE=0.30, *p*=0.008; 95% *CI* [0.21, 1.41], *d = 0.71).* BOLD response was also found to be significantly decreased in the Fear>Neutral+Happy (*M_diff_*=-0.67, SE=0.30, *p*=0.029; 95% *CI* [0.07, 1.27], *d = 0.62*) and increased in the Happy>Neutral+Fear (*M_diff_*=0.67, SE=0.30, *p*=0.028; 95% *CI* [0.07, 1.27], *d = 0.6*) conditions. No significant changes were observed one-month after the 1 mg control dose or in the right amygdala following either dose (all *p*>0.05).

**4.4.	Figure S10. Left amygdala BOLD response to emotional faces**

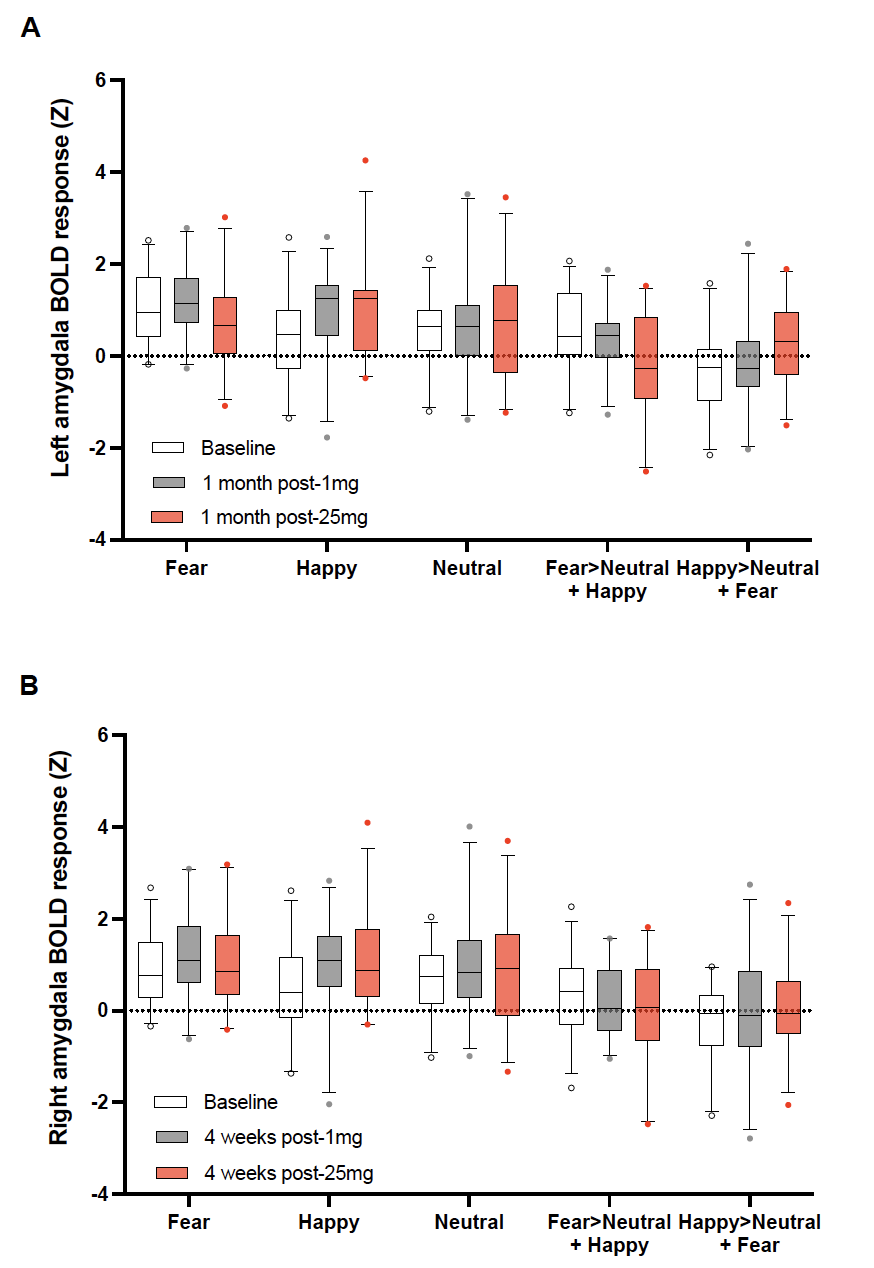

**Figure S10. Box and whisker plot for the left amygdala response to emotional faces.** From the Harvard-Oxford atlas, we threshold at 50% probability and then extract the values from the entire left amygdala, given that threshold. We elected not to run statistical significance tests on any paired contrasts.

**4.5.	Figure S11. Right amygdala BOLD response to emotional faces**

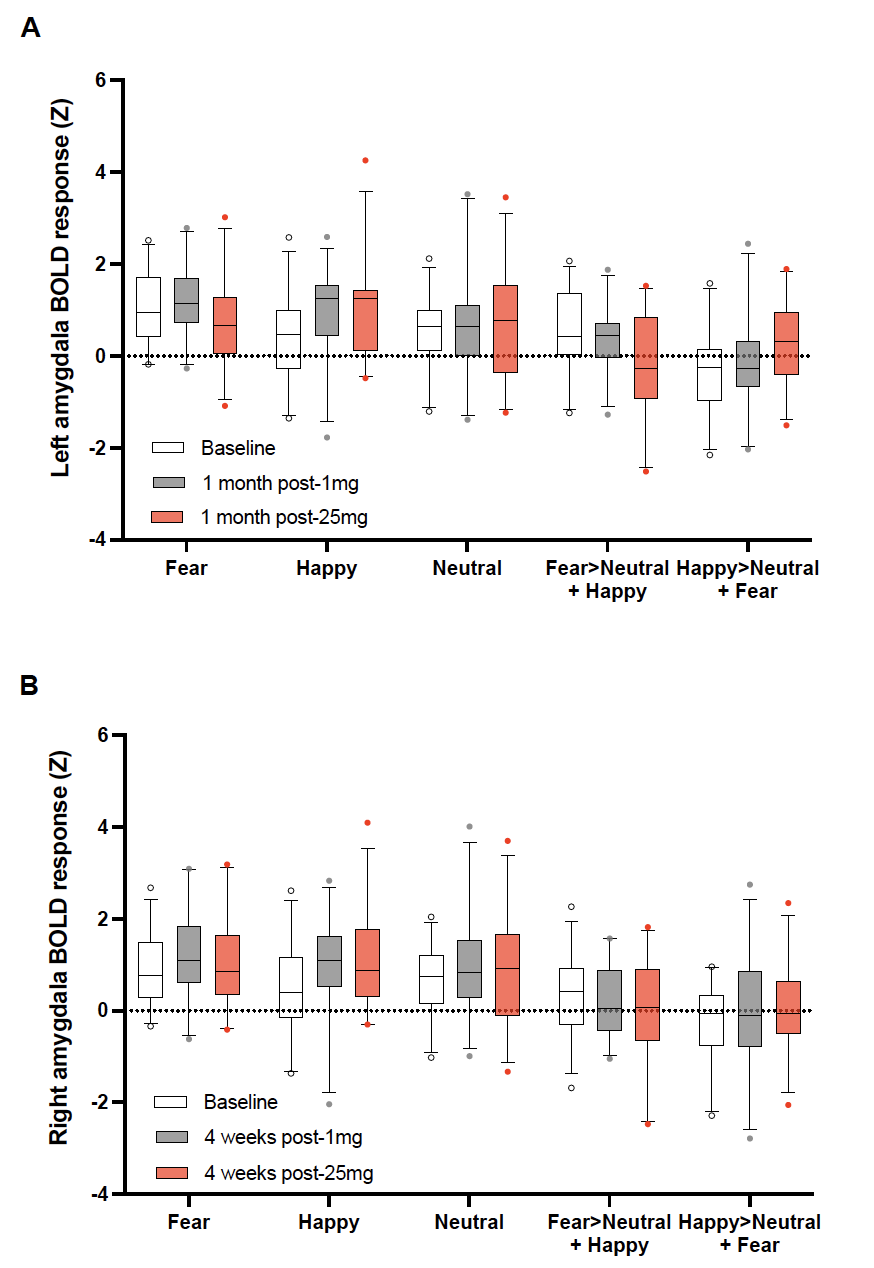

**Figure S11. Box and whisker plot for the right amygdala response to emotional faces.** Unlike for the left amygdala, no contrasts were statistically significant for the right amygdala. From the Harvard-Oxford atlas, we threshold at 50% probability and then extract the values from the entire right amygdala, given that threshold.

**4.6.	Table S12. Effects of high-dose psilocybin on amygdala functional connectivity (general psychophysiological interaction, gPPI)**

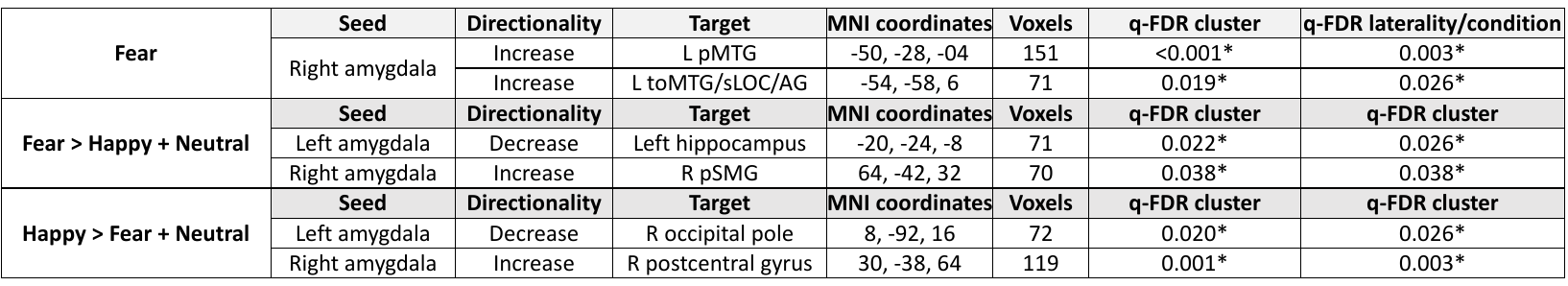

**Table S12. Effects of high-dose psilocybin on amygdala functional connectivity.** Time contrast: post-1mg > post-25mg; L=left, R=right; pMTG = posterior middle temporal gyrus; toMTG = temporo-occpital MTG; sLOC = superior lateral occipital cortex; AG = angular gyrus; pSMG = posterior supramarginal gyrus; * = q-FDR<0.05.

Psychophysiological interaction results in the face paradigm suggest emotion-specific modulation of amygdala circuitry. In the fear faces condition, amygdala functional connectivity was increased between the amygdala and temporal, parietal, and occipital areas, while amygdala-hippocampus coupling decreased. In the happy faces condition, functional connectivity was increased between the amygdala and the postcentral gyrus, while amygdala-occipital pole functional connectivity was decreased. These results are broadly consistent with a prior PPI amygdala analysis involving scanning after psilocybin-therapy for treatment-resistant depression ^1^, but also highlight amygdala-temporal lobe and amygdala-hippocampal functional connectivity changes that align with some previous work in depression ^2,3^. However, some difference between this present study’s results and prior work ^1^ were also apparent, such as a lack of decreased ventromedial PFC-amygdala functional connectivity.

**4.7. Figure S13.** **Effects of high-dose psilocybin on amygdala functional connectivity**

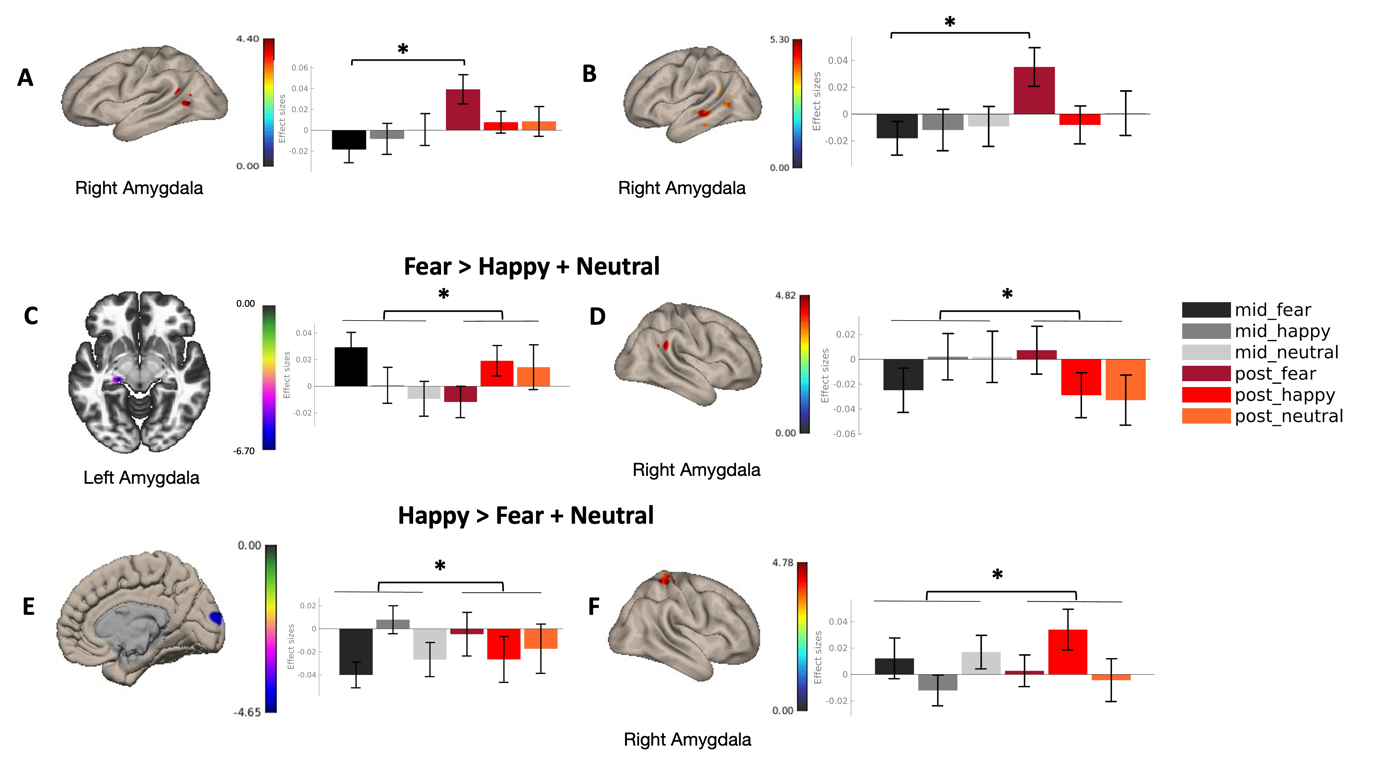

**Figure S13**. A-B = fear-25mg > fear-1mg; C-D = (fear-25mg > happy-25mg + neutral-25mg) > (fear-1mg > happy-1mg + neutral-1mg); E-F = (happy-25mg > fear-25mg + neutral-25mg) > (happy-1mg > fear-1mg + neutral-1mg); A = R amygdala-posterior middle temporal gyrus, B = R amygdala-temporal occipital middle temporal gyrus, C = L amygdala-hippocampus, D = R amygdala-posterior supramarginal gyrus, E = L amygdala-occipital pole, F = R amygdala-postcentral gyrus.

**4.8. Figure S14.** **Bilateral amygdala BOLD RSFC analysis.**

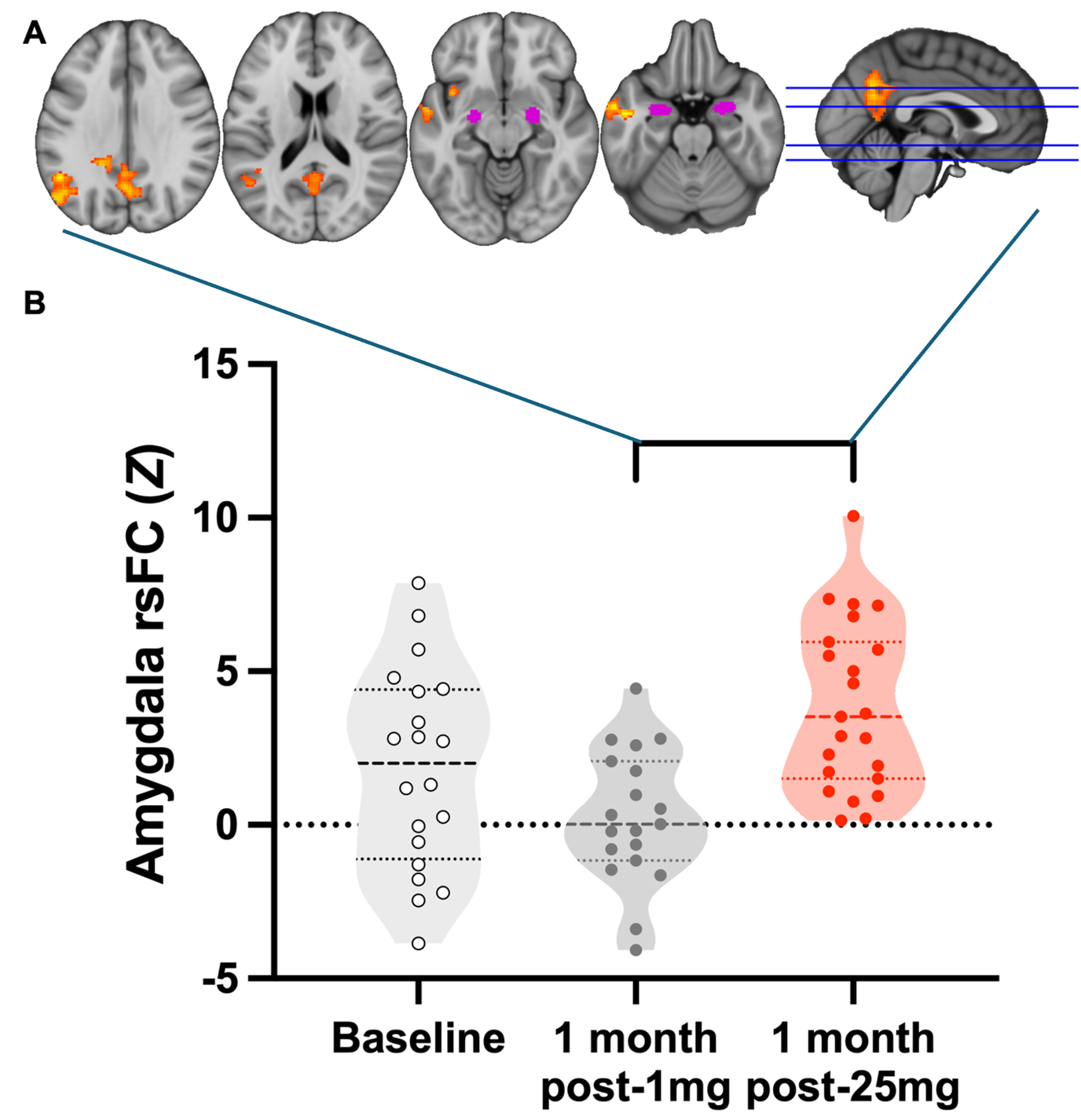

**Figure S14. A)** The bilateral amygdala is shown in magenta, from the Harvard-Oxford atlas, >50% threshold. Hot colours show the cluster-corrected map (threshold Z = 2.3, p < 0.05) for amygdala RSFC contrast, one-month post-25mg vs one-month post-1mg. Increased amygdala coupling can be seen with regions that overlap the so-called ‘default-mode network’. Values from this timepoint-specific contrast, and the clusters therein, contributed to the amg-RSFC correlation matrix shown in Figure 4B of the main paper. **B)** Violin plot of bilateral amygdala BOLD RSFC analysis. Coupling strength (Z stat) values are derived from a mask of the positive result for the contrast one-month post 25mg vs 1 month post 1mg, as shown in **A**. We refrain from statistical tests on these contrast-selective voxels from which the values in **B** derive. Note that this analysis is repeated as an ANOVA across all timepoints in **Figure S16, below.**

**4.9. Figure S15. Amygdala RSFC with single subject values shown.**

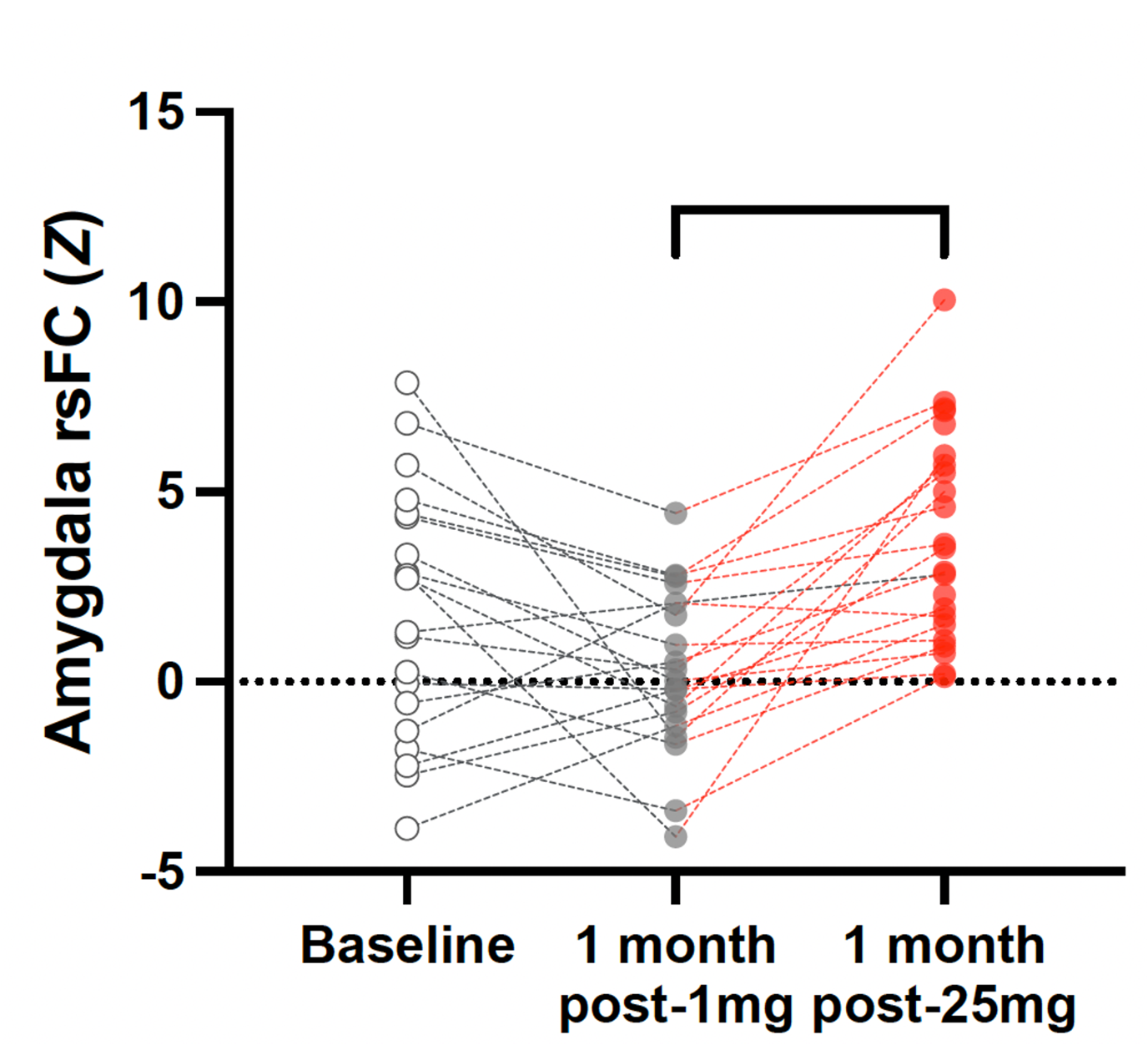

**Figure S15. Amygdala RSFC with values from the 25mg vs 1mg contrast.** This figure is intended for descriptive purposes only, to show the single subject values. Note that values derive from the cluster that was significant in the one-month post 1mg versus one-month post 25mg contrast (i.e., shown in Fig S14). Note that this analysis is repeated as an ANOVA across all timepoints in Figure S16, below.

**4.10. Figure S16. Whole-brain naïve ANOVA across all timepoints**

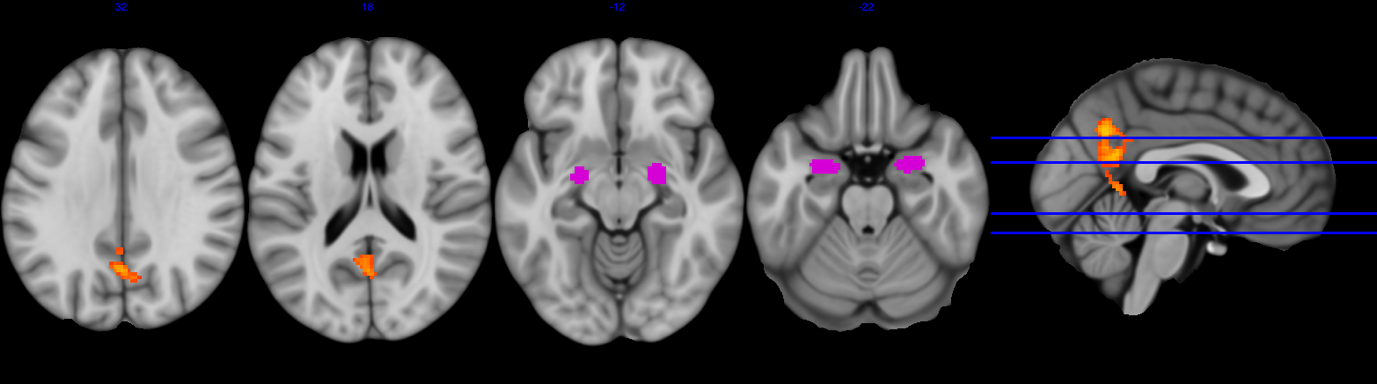

**Figure S16. Bilateral amygdala RSFC analysis as ANOVA across all timepoints.** The above map shows different views of the bilateral amygdala seed (magenta) in a RSFC analysis across all three timepoints, where the cluster (z = 2.3, cluster corrected, p < 0.05) was sensitive to *time*.

**4.11. Validation analysis: in-scanner anxiety: amygdala RSFC analysis**

For the amygdala ROI rsFC analysis, we examined in-scanner visual analogue scale (VAS) ratings completed immediately after each resting state run revealed a significant effect of anxiety (F(1.16,28.42)=7.08, p=0.010), with significantly greater anxiety among the participants at pre-intervention baseline relative to one month after 1 mg psilocybin; Mdiff=1.00, SE=0.40, p=0.013; 95% CI [0.18, 1.82], d=0.7) and one month after 25mg; Mdiff=1.14, SE=0.33, p=0.010; 95% CI [0.33, 2.19], d=0.7). When anxiety levels during the baseline scan were entered into regression models as a covariate, the one-month post-1mg versus baseline scan changes in amygdala rsFC no longer reached significance (F(1,19)=1.35, p=0.259 ns) – implying that any initial observations of change in amygdala RSFC post 1mg may have been related to unusually high anxiety levels for the baseline scan (in this resting-state scan only), rather than e.g., a direct effect of 1mg psilocybin itself.

Further strengthening this inference on the role of in-scanner anxiety in the baseline scan as a confounding variable, a correlation was found between the drop in anxiety in the one-month post-1mg scan (vs the baseline scan) and parallel changes in amygdala rsFC (R2=0.206, p=0.026). Outlier analyses revealed three outliers with especially high anxiety ratings (i.e., a VAS score of >60%) for their baseline scan. When these were removed, and the analyses repeated, Bonferroni-corrected post-hoc comparisons showed a robust increase in amygdala RSFC one month after 25mg psilocybin (Mdiff=3.42, SE=0.65, p<0.0001; 95% CI [1.78, 5.01], d=1.2), and no changes one-month after the 1mg dose. We therefore decided to use this result for the main amygdala RSFC analyses shown here in this supplement (e.g., Figure S14-16); these are also the data used in the correlation plot in the main paper.

**4.12. Resting State Networks (RSN) – Within and Between Networks RSFC**

RSNs were derived using Independent Component Analysis (ICA) performed on Human Connectome Project data (Van Essen et al., 2013). In summary, 20 independent components (ICs) were derived, of which the same 12 functionally meaningful RSNs were identified, namely: medial visual network (VisM), lateral visual network (VisL), occipital pole network (VisO), auditory network (AUD), sensorimotor network (SMN), default-mode network (DMN), parietal cortex network (PAR), dorsal attention network (DAN), salience network (SAL), posterior opercular network (POP), left frontoparietal network (lFPN) and right frontoparietal network (rFPN). (See Roseman et al. (2014) for more details).

Network integrity (Within-RSN rsFC) was calculated for each RSN for both pre-treatment and post-treatment. All 20 HCP ICA components were entered into FSL’s dual regression analysis (Beckmann et al., 2009). The first step of the dual regression used the components as regressors applied to the 4D BOLD datasets for each subject, resulting in a matrix of time-series for each ICA. The second step involved regressing these time-series into the same 4D scan data to get a subject-specific set of spatial maps (parameter estimate (PE) images). For each subject and for each condition, within each of the 12 RSNs of interest (threshold = 3), the mean PE across voxels was calculated. This mean PE represents the integrity value. Subsequently, paired t-tests were used to calculate the difference in integrity between conditions for each RSN (Bonferroni corrected for 12 RSNs).

Between-RSN rsFC was calculated in a similar manner to previous analyses involving acute LSD (Carhart-Harris et al., 2016) and psilocybin (Roseman et., 2014). Specifically, a 12 × 12 matrix was constructed representing rsFC between different RSN pairs. For each subject and for each condition, the time-series for the relevant pair of RSNs, was entered into a GLM, resulting in a PE value representing the strength of functional connectivity between them. GLM was used rather than correlation coefficients because differences between Pearson’s correlations could be a result of either signal or noise differences; therefore, it is preferable to perform regression and look for differences on the PE (Friston, 2011). The GLM was estimated twice: 1) each RSN as a dependant variable in one model, and 2) each RSN as an independent variable in the second model. These two PE values were then averaged together, to generate a symmetric 12 × 12 matrix. Paired t-tests (two-tailed) were used to compare the PE values of scan 2 and scan 3. There were no significant changes for both Within-RSN RSFC, and Between-RSN RSFC.

**4.13. Figure S17. Modularity with single subject values shown.**

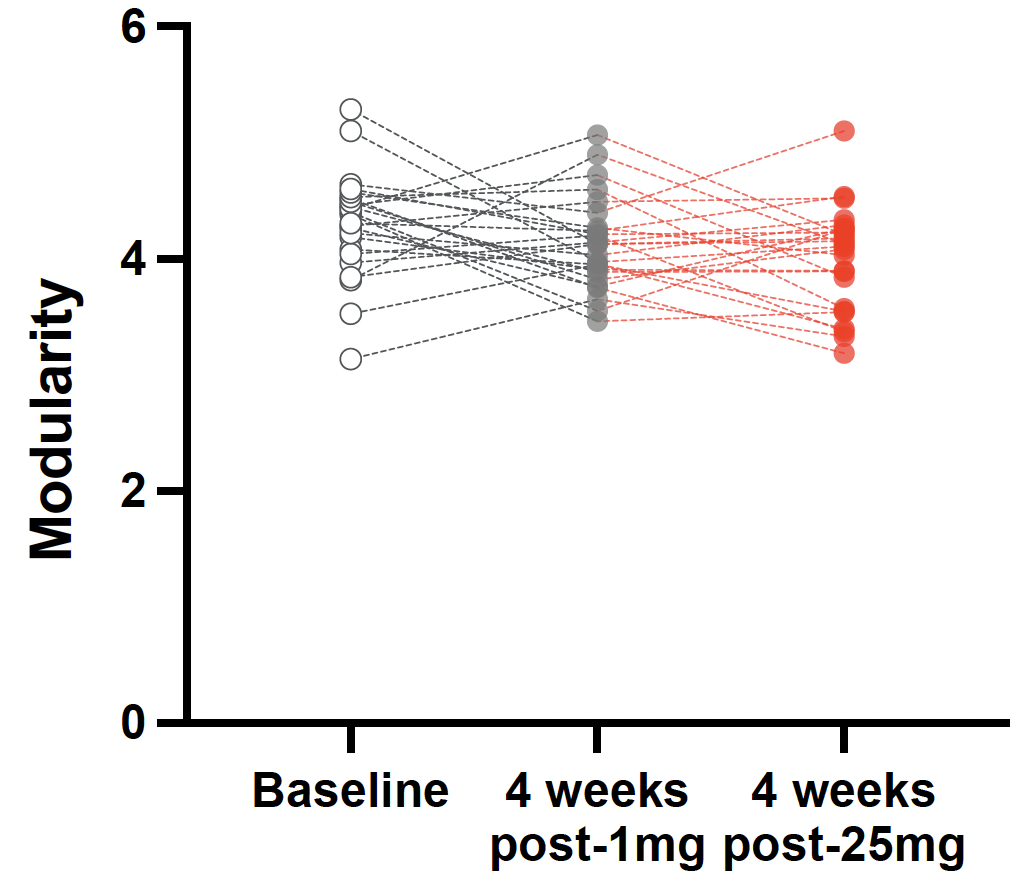

**Figure S17.** Modularity with single subject values shown and lines connecting all timepoints. There was no significant effect of time on brain network modularity.

5. **DTI outcomes**

**5.1. Figure S18. DTI: tracts where changes were observed.**

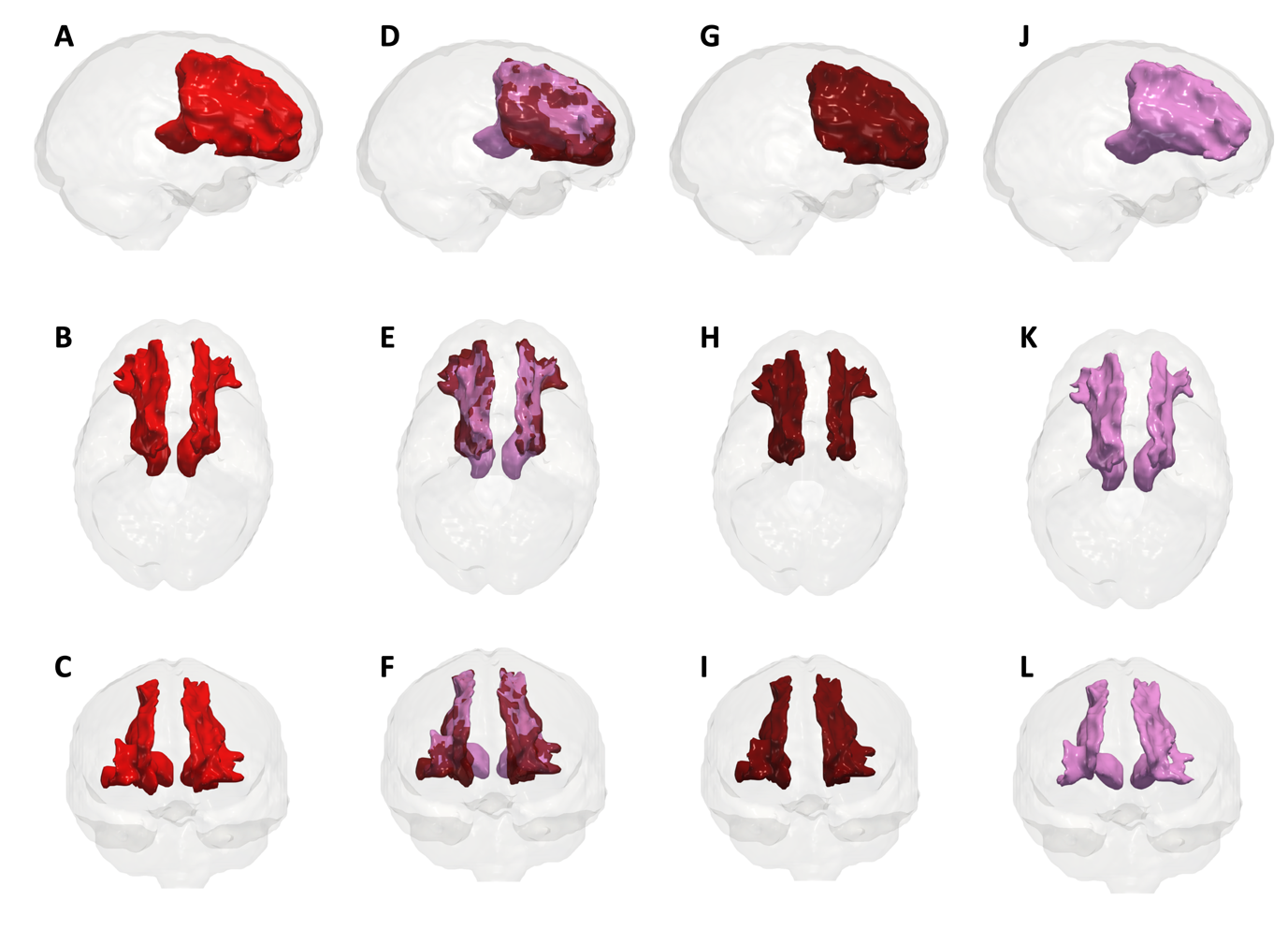

**Figure S18. Prefrontal cortex to subcortical tracts:** PFC = prefrontal cortex; THA = thalamus; STR = striatum. A-C) Combined PFC-THA & PFC-STR tracts; D-F) Combined PFC-THA (pink) & PFC-STR (dark red) tracts; G-I) PFC-STR tracts - alone; J-L) PFC-THA tracts - alone. Top-to-bottom, sagittal, axial, and coronal planes. All tracts where there was a significant decrease in axial diffusivity one-month after 25mg psilocybin.

**5.2.	Figure S19. Axial diffusivity values. Left hemisphere.**

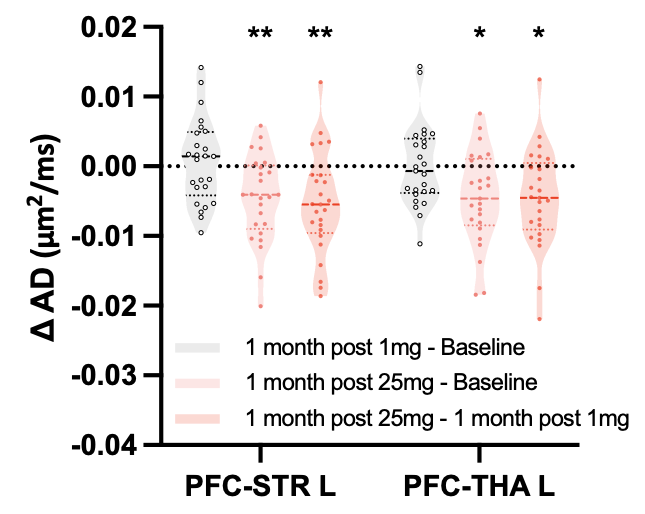

**Figure S19. Axial diffusivity values without free-water correction. Left hemisphere.** Post-hoc t-tests revealed a significant decrease in AD one-month post-25mg vs. one-month post-1mg in PFC-STR (t(24) = -3.09, p = 0.030) and PFC-THA (t(24) = -3.83, p = 0.005). Change in mean axial diffusivity (AD) in the left prefrontal-thamalus (PFC-THA) tract without free-water correction is shown in the violin plot above. The change in the right hemisphere was in the same direction but was not statistically significant after correction for multiple comparisons. * = p < 0.05, ** = p < 0.01, corrected for multiple comparisons.

**5.3. Figure S20. Axial diffusivity values with single subject datapoints shown.**

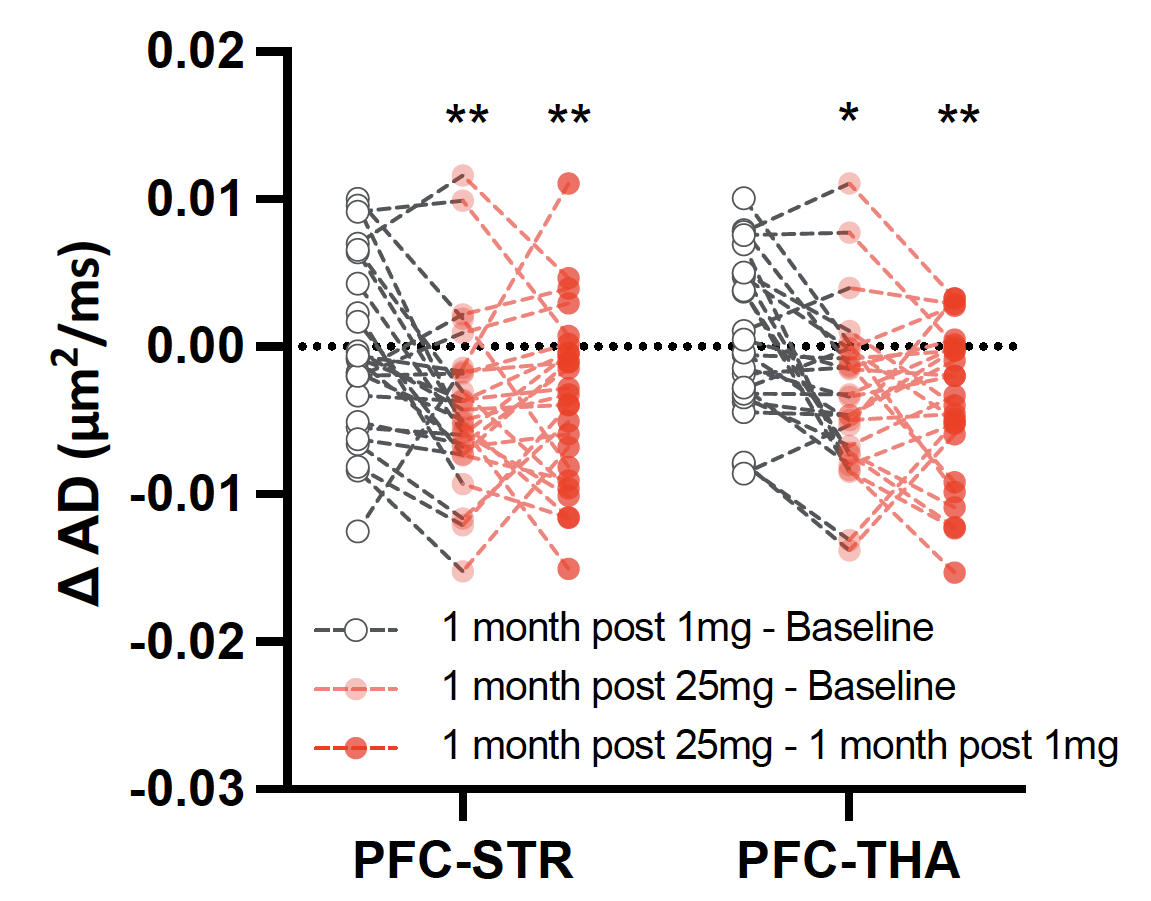

**Figure S20.** Axial diffusivity but with single subject datapoints shown (without free-water correction, hemispheres merged). Note that all datapoints are deltas i.e., subtractions of one timepoint from another. No changes were observed after the 1mg psilocybin (white circles). Changes only appeared after the 25mg dose of psilocybin (faint red and full red circles). * p < 0.05, ** p < 0.01.

**5.4. Figure S21. Fractional Anisotropy (FA) with free-water correction.**

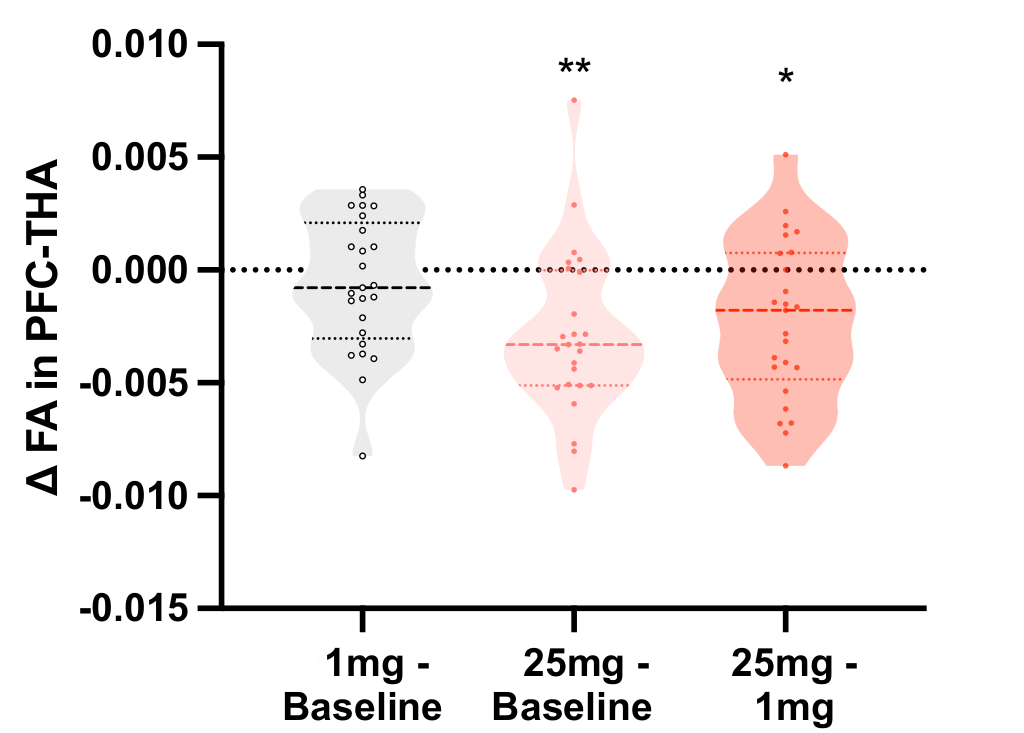

**Figure S21.** Change (decreases) in mean free-water corrected fractional anisotropy (FA) in the bilateral prefrontal-thalamus (PFC-THA) tracts one-month post-25mg. Only post-25mg contrasts were significant, again implying that the DTI change was dependent on the high-dose of psilocybin. As with the other DTI metrics, there was no change after the 1mg dose (leftmost violin in grey). * = p < 0.05, ** = p < 0.01, corrected for multiple comparisons.

ANOVAs on Free-water-corrected DTI revealed a significant effect of time on FA in the PFC-STR tract (F(2, 48) = 7.88, p = 0.026) and PFC-THA tract (F(2, 48) = 9.98, p = 0.006), as well as the genu of the corpus callosum (F(2, 48) = 7.09, p = 0.048). Post-hoc t-tests with revealed a significant decrease in FA one-month post-25mg vs. one-month post-1mg in PFC-THA (t(24) = -3.19, p = 0.035).

**5.5. Free-water correction values for axial diffusivity**

We observed a consistent change in AD with free-water-corrected DTI. ANOVA revealed a *Time* x *Dose* interaction on AD in PFC-STR (F(2, 48) = 10.27, p = 0.005) and PFC-THA (F(2, 48) = 12.79, p = 0.001

6. **Psychological outcomes**

**6.1. Detailed description of Intradimensional/extradimensional (IDED) task**

The IDED task is comprised of 9 phases, which are detailed below:

1. Simple discrimination: Basic trial and error requiring participants to discriminate between one of two stimuli in the same dimension (yellow shapes). This phase acts to determine the initially correct stimulus.
2. Simple reversal: Reversal of correct stimulus, whereby previously identified stimulus will now prompt a failure notification.
3. Compound discrimination with second dimension: The addition of a second compound dimension, a distracting element (blue lines). The correct response remains the pre-existing stimulus (yellow shape), and therefore the new element must be ignored.
4. Compound discrimination: Two dimensions (yellow shapes and blue lines) become overlapped, with correct stimulus of the pair (yellow shapes) unchanged.
5. Compound reversal: Correct choice remains in the same dimension (yellow shapes), but moves to the other element of the pair.
6. Intra-dimensional shift: The illustrative stimuli change, but the correct response remains within the same dimension (yellow shape). This phase acts as a specific measure of attentional set-shifting.
7. Intra-dimensional reversal: Correct choice becomes the second element of the pair within the same dimension (yellow shapes).
8. Extra-dimensional shift: The illustrative stimuli change again, with the correct stimulus now being within the opposite dimension (blue lines). This phase examines both attentional set-shifting and reversal learning.
9. Extra-dimensional reversal: The correct stimulus swaps to the opposite element of the pair in the same dimension (blue lines).

**6.2. Figure S22. IDED task phase 1-9.**

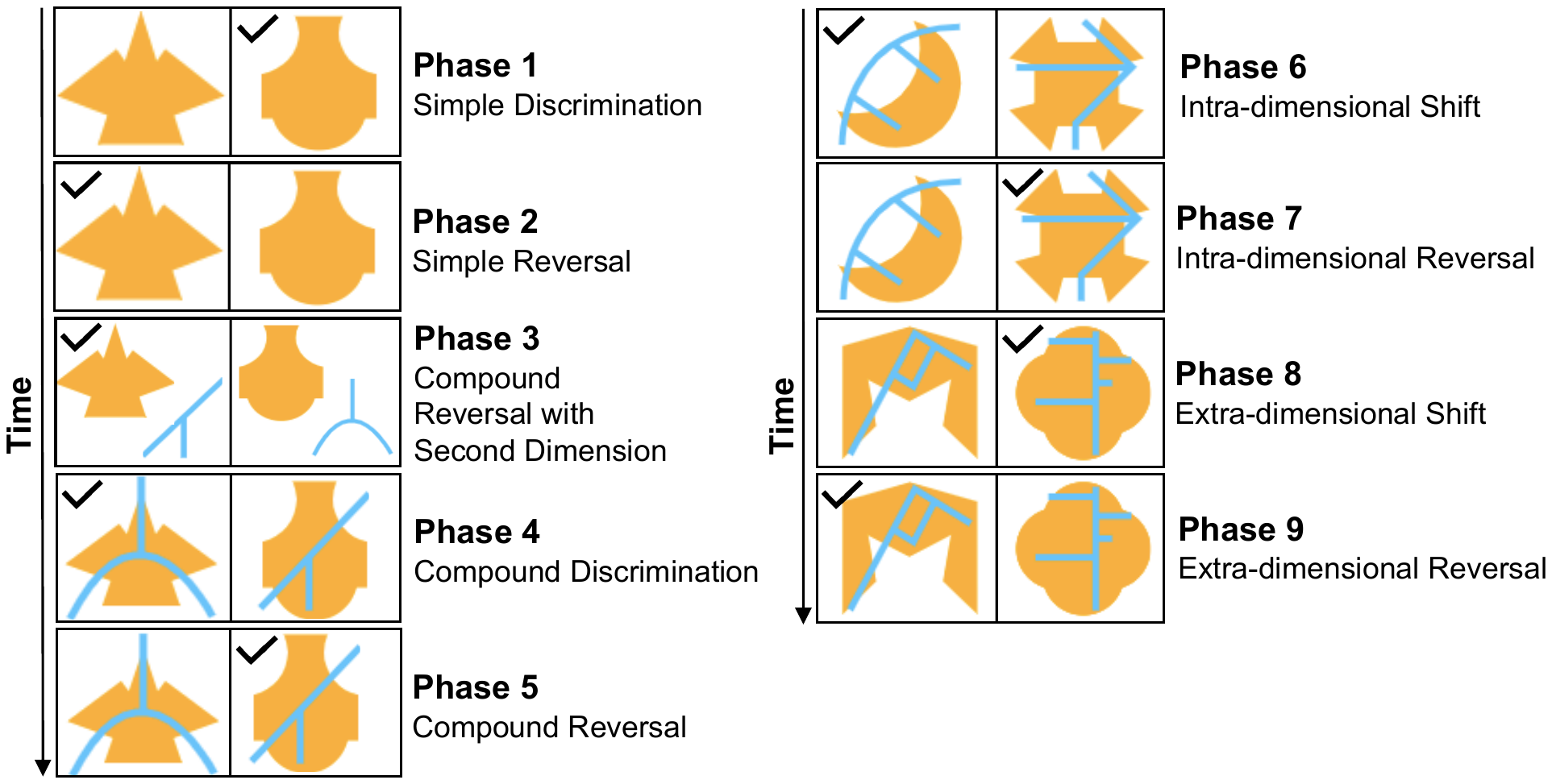

**6.3. Figure S23. IDED. Cognitive flexibility (more granular) findings.**

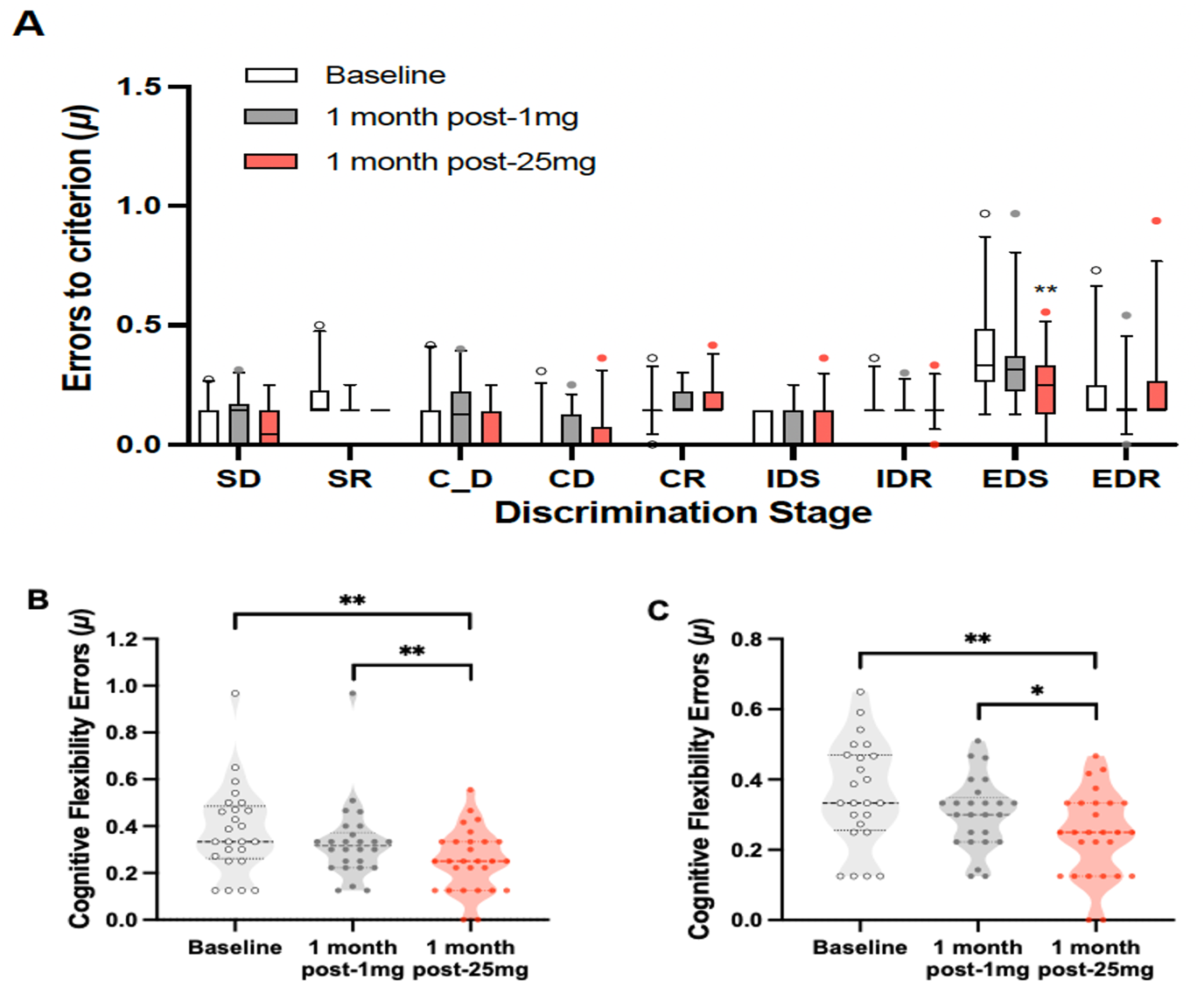

**Figure S23.** **A**) The IDED with its various sub-parameters shown. Abbreviations: Intra-dimensional/extra-dimensional task; SD, simple discrimination; C_D, compound discrimination with second dimension; CD, compound discrimination; IDS, intra-dimensional shift; EDS, extra-dimensional shift: SR, simple reversal; CR, compound reversal, IDR, intra-dimensional reversal; EDR, extra-dimensional reversal. EDS is considered the most relevant index of cognitive flexibility; this is the parameter we measure and refer to as “cognitive flexibility errors” in B & C. ** = p < 0.01. **B**) EDS with all data included, including an outlier participant (*M_diff_*=0.13, SE=0.04, *p*=0.008; 95% *CI* [0.02, 0.12], *d*=0.6). ** = p < 0.01. **C**) We also conducted an outlier analysis using the ROUT method (for ‘**R**obust regression and **Out**lier removal’) which identified one significant outlier across two timepoints (i.e., pre-dose baseline and one-month post-1mg). Figure C therefore shows the results with this outlier removed. As this outlier scored in the direction of our findings, we report C in the main manuscript (*M_diff_*=0.06, SE=0.02, *p*=0.016; 95% *CI* [0.00, 0.12], *d*=0.5), i.e., we report the more conservative analysis in the main paper and show here how the main result **was not** driven by the outlier. * = p < 0.05, ** = p < 0.01.

**6.4. Figure S24. EDS errors. Single subject data**

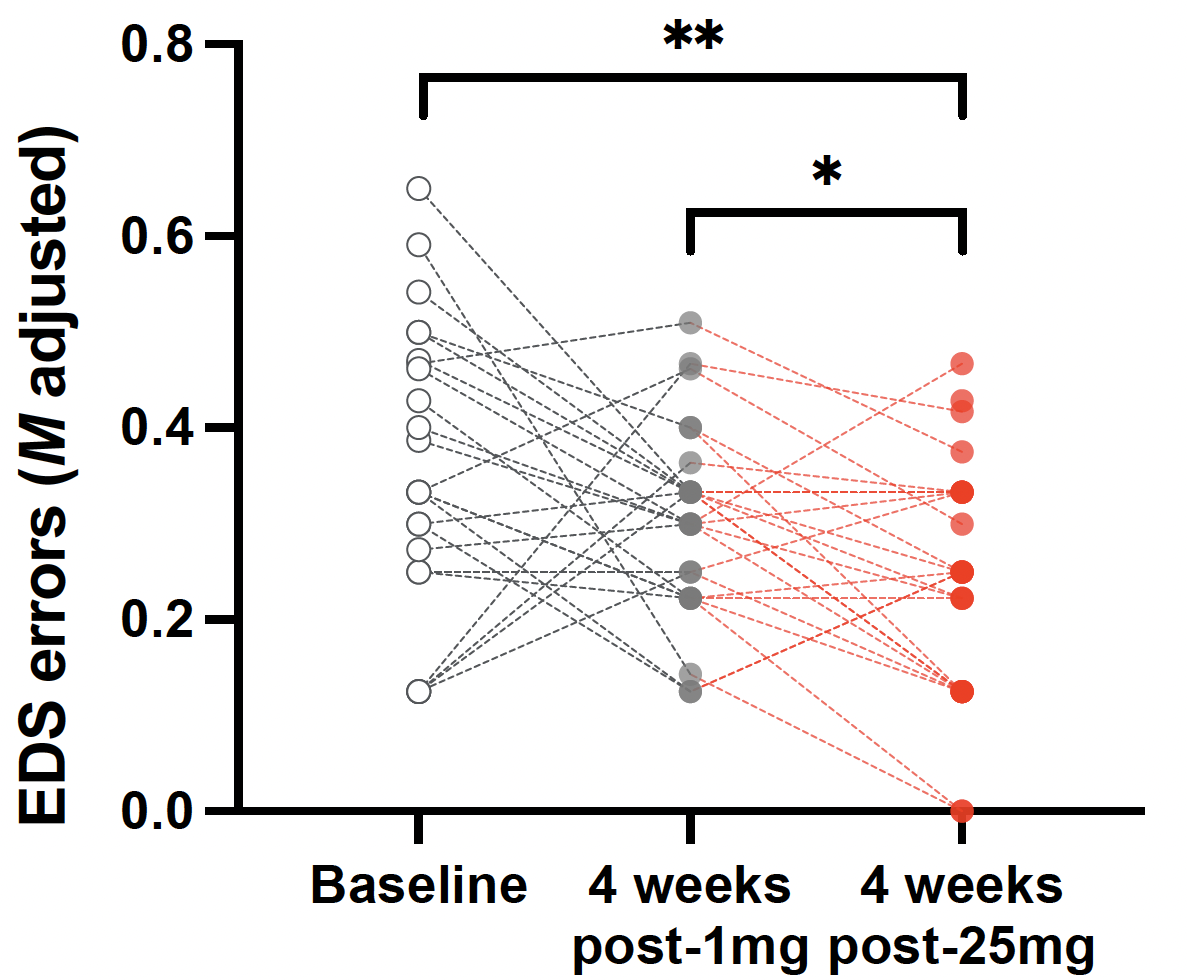

**Figure S24.** EDS errors showing single subject scores. Asterisks show significance at the group mean level, i.e., * p< 0.05, ** p< 0.01.

**6.5. Table S25. IDED task errors per criterion**

| **IDED  task stages** | | **Baseline** | **1 mg + 4 weeks** | **25 mg + 4 weeks** |
| --- | --- | --- | --- | --- |
|  |  | Adjusted Errors (μ)*  μ ± *SEM* | | |
| *Discrimination* | **SD** | 0.099 ± 0.027 | 0.111 ± 0.021 | 0.076 ± 0.016 |
|  | **C_D** | 0.137 ± 0.036 | 0.126 ± 0.027 | 0.079 ± 0.018 |
|  | **CD** | 0.034 ± 0.015 | 0.045 ± 0.014 | 0.045 ± 0.017 |
|  | **IDS** | 0.068 ± 0.014 | 0.070 ± 0.016 | 0.076 ± 0.018 |
|  | **EDS** | 0.383 ± 0.038 | 0.329 ± 0.032 | 0.254 ± 0.025 |
| *Reversal* | **SR** | 0.195 ± 0.018 | 0.155 ± 0.007 | 0.143 ± 0.000 |
|  | **CR** | 0.156 ± 0.012 | 0.172± 0.010 | 0.183 ± 0.013 |
|  | **IDR** | 0.160 ± 0.010 | 0.156 ± 0.007 | 0.154 ± 0.010 |
|  | **EDR** | 0.217 ± 0.028 | 0.164 ± 0.018 | 0.248 ± 0.035 |

**Table S25.** Values for the IDED. All values are mean errors per criterion, adjusted for learning rate. Abbreviations: IDED, Intra-dimensional/extra-dimensional task; SD, simple discrimination; C_D, compound discrimination with second dimension; CD, compound discrimination; IDS, intra-dimensional shift; EDS, extra-dimensional shift: SR, simple reversal; CR, compound reversal, IDR, intra-dimensional reversal; EDR, extra-dimensional reversal.

**6.6. Figures S26. Adapted version of ‘Persisting Effects Questionnaire’ of Griffiths et al. 2006.**

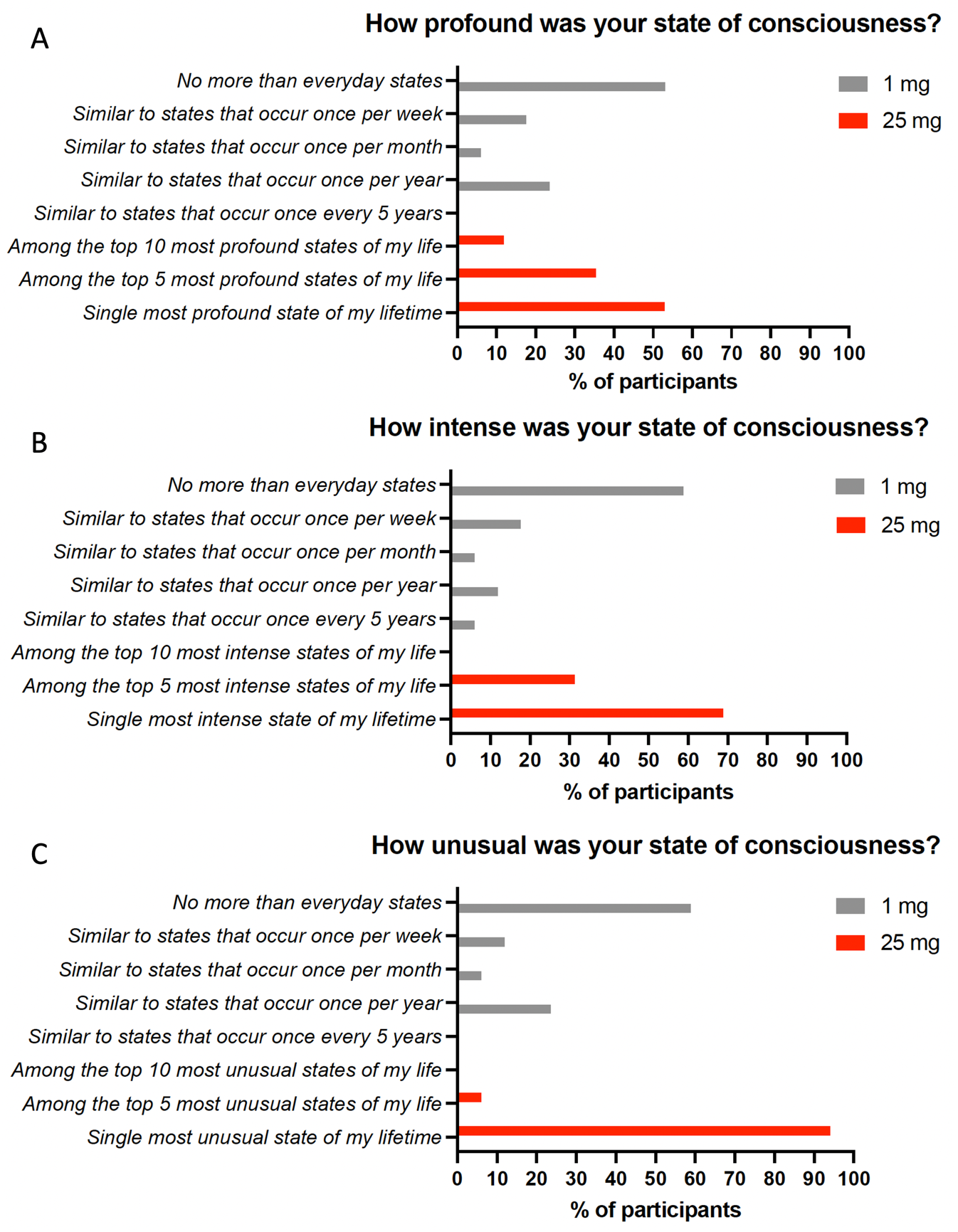

**Figure S26.** One-month after each dosing session, we asked all participants to assess the quality of the state of consciousness they had experienced during each dosing session using the labelled ranking criteria shown on the y-axis of the above charts. Participants ranked how: **A)** *profound*, **B**) *intense*, and **C**) *unusual* their state of consciousness was in relation to their life up to that moment. They did this after both the 1mg (gray) and 25mg psilocybin experiences (red). This scale was an adapted version of the Persisting Effects Questionnaire used in ^4^.

7. **References**

1	Mertens, L. J. *et al.* Therapeutic mechanisms of psilocybin: Changes in amygdala and prefrontal functional connectivity during emotional processing after psilocybin for treatment-resistant depression. *J Psychopharmacol* **34**, 167-180 (2020). <https://doi.org/10.1177/0269881119895520>

2	Wackerhagen, C. *et al.* Amygdala functional connectivity in major depression - disentangling markers of pathology, risk and resilience. *Psychol Med* **50**, 2740-2750 (2020). <https://doi.org/10.1017/S0033291719002885>

3	Tang, S. *et al.* Abnormal amygdala resting-state functional connectivity in adults and adolescents with major depressive disorder: A comparative meta-analysis. *EBioMedicine* **36**, 436-445 (2018). <https://doi.org/10.1016/j.ebiom.2018.09.010>

4	Griffiths, R. R., Richards, W. A., McCann, U. & Jesse, R. Psilocybin can occasion mystical-type experiences having substantial and sustained personal meaning and spiritual significance. *Psychopharmacology* **187**, 268-283; discussion 284-292 (2006). <https://doi.org/10.1007/s00213-006-0457-5>
