## Supplementary material for "Human brain changes after first psilocybin use": Methods section

**Materials and methods**

**RESOURCE AVAILABILITY**

**Lead contact.**

Further information and requests should be directed to Taylor Lyons or Robin Carhart-Harris.

**Materials availability.**

No unique reagents were generated for this study.

**Data and code availability.**

Information required to reanalyze the data reported in this paper is available from the lead contacts upon request.

**EXPERIMENTAL MODEL AND SUBJECT DETAILS**

**Ethical approvals and drug procurement.**

This study was approved by the London-Surrey Research Ethics Committee and sponsored by the Joint Research and Compliance Office, Imperial College London. The National Institute for Health Research/Wellcome Trust Imperial Clinical Research Facility (ICRF) provided site-specific approvals. This research was carried out in accordance with Good Clinical Practice guidelines. All participants provided written informed consent.

An algorithm provided by the UK Medicines and Healthcare products Regulatory Agency (MHRA) was followed to assess whether the present study should be designated as a clinical or non-clinical trial. The algorithm and the MHRA confirmed that the present study was not a clinical trial. This was further confirmed by the sponsoring institution (Imperial College London) in the process of their sponsorship. Their decisions justify the designation of this study as an exploratory, translational study in healthy volunteers. In accordance with this designation, the study was not pre-registered as a clinical trial.

Bottled and encapsulated (size 4) psilocybin was provided by COMPASS Pathways and securely stored in Imperial College London in light-protected and temperature-controlled conditions. A Home Office Schedule 1 Drug License for the storage and handling of psilocybin was obtained. Psychotherapeutic and capsule (size and color matched) ingestion procedures were consistent across dosing days.

**Study design and participants.**

This controlled, fixed-order, within-subjects study investigated the effects of psilocybin in healthy human adults with no prior psychedelic experience (*N*=28). All participants received two oral doses of psilocybin, 4-weeks apart: (1) a control dose of 1mg psilocybin on the first dosing day, considered to be a subthreshold dose that is unable to occasion a psychedelic experience; and (2) a fully active dose of 25mg psilocybin, considered to be a high dose and capable of inducing profound psychedelic effects, 4-weeks or 1-month later. This fixed order design was necessary given the hypothesized carry-over effects of 25mg psilocybin. To uphold blinding and control for expectancy effects, participants were informed that they would receive psilocybin on both sessions of a variable dose up to 25mg. No further information regarding dosage was provided. For a schematic of the design and timeline of interventions, see Figure S1 and Table S2.

Participants (N=28) had an average age of 41 years (*SD=*8.7, range: 29–59) and were balanced in terms of gender (*χ^2^*=0.57, *p=*0.450) and educational attainment (*χ^2^*=0.62, *p=*0.430). All participants were naïve to psychedelic drugs and the majority were British (75%; *χ^2^*=7.00, *p<*0.01) and Caucasian (86%; *χ^2^*=14.57, *p<*0.001) in full-time employment (86%; *χ^2^*=14.29, *p<*0.001). A breakdown of the participant demographic details can be found in Table S3. Recruitment information and full inclusion and exclusion criteria are detailed in the supplementary file.

**METHODS DETAILS**

**Acute dosing procedures.**

Participants refrained from caffeinated products and consumed a very light breakfast >1 hour before arrival on dosing days. Upon arrival, participants were breathalyzed and provided a urine sample for drugs of abuse and pregnancy (where applicable). Participants set an intention for the psilocybin sessions before dosing, which was written on a white board in the dosing room. Dosing took place in a dimly lit, aesthetically pleasing room within the ICRF facility. A music playlist was created on Spotify (<https://open.spotify.com/playlist/7xcb0s46JhffYOY8euOEFO?si=1dc0dd49ac8a44dd>) and simultaneously played through both headphones (Bose Sound-link II around ear) and speakers (KRK.RP5G3 Rockit Powered Studio). Participants wore a MindFold eye mask and were encouraged to lie in supine position on the bed. Both dosing days involved non-directive, guided supervision throughout. At the end of dosing days, participants were assessed for potential discharge by the study physician.

**EEG procedure*.***

To assess acute brain effects, a 24-channel wireless EEG cap (DSI-24 System, Wearable Sensing) with 21 dry electrodes (0.317 µV resolution, 300 Hz sampling rate and with an integrated amplifier, was used on dosing days. Electrodes were positioned following the 10-20 international format: Fp1, Fp2, Fz, F3, F4, F7, F8, Cz, C3, C4, T7/T3, T8/T4, Pz, P3, P4, P7/T5, P8/T6, O1, O2. Electrode Pz served as an online reference, with data re-referenced to electrodes A1 and A2 placed on the earlobes for the subtraction of noise in other channels. Fpz was selected as the ground electrode. All data were recorded using a Bluetooth-connected DSI-Streamer-v.1.08.41.

Four eyes-closed resting-state EEG (rsEEG) recordings, each lasting 4 minutes, were completed in silence at baseline (pre-dosing) and at ~1 hour, ~2 hours, and ~4.5 hours post-dosing. All EEG data were preprocessed using the Fieldtrip toolbox^1^ in MATLAB (R2019B, MathWorks, Inc). rsEEG data were band-passed filtered at 1-45 Hz and visually inspected for gross artifacts. Noisy segments and channels were removed and interpolated, respectively. Independent Component Analysis (ICA) was used for the removal of eye blink artifacts. Next, data was re-referenced to the average and segmented in 2 second-epochs. Following preprocessing, spectral analysis was performed using Slepian multitapers with spectral smoothing of +/- 0.5 Hz using the Fieldtrip toolbox^1^. Resulting spectra were divided in the following frequency bands for statistical analysis: *Delta* (1-4 Hz), *theta* (4-8 Hz), *alpha* (8-13 Hz), *beta* (13-30 Hz), and *gamma* (30-45 Hz). Scalp-level analysis of peak effects of 25mg on EEG measures were performed using paired-*t*-tests at each electrode. Multiple comparisons of EEG results were controlled using cluster randomization analysis with an initial cluster-forming threshold of *p*=0.05 repeated for 5000 permutations.

**EEG LZc models**

To investigate if acute brain effects were associated with subsequent cognitive and behavioral changes, a procedure was designed to build a predictive model of a given outcome based on the values of a predictor at different electrodes and timepoints. In particular, our conjecture was that the dynamical trajectory of LZc could predict the level of psychological insight rated by participants via the PIS the day after dosing. The results shown in Figures 4A and C are relevant in this regard. Figure 4D uses values across *all* sensors but at the 2h timepoint when we know the intensity of subjective effects and plasma concentrations of psilocybin’s active metabolite, psilocin, are maximal. ^2^

Data-driven analyses using acute rsEEG LZc for predicting psychological outcomes (insight and well-being changes) were carried out in a four-step procedure:

1) First, correlations between the target variable and LZc values across subjects were computed for each electrode and for each timepoint separately. All electrode-timepoint combinations were ranked according to their correlation with the target variable.

2) Next, we constructed a predictor based on the average LZc of all electrode-timepoints whose correlation to the target variable was above a given value ϑ. The performance of this predictor was assessed by the out-of-sample R^2^, computed via leave-one-out cross-validation. Specifically, we used *n*-1 subjects to (i) find the electrode-timepoint combinations that exceed the threshold, (ii) take the average LZc of each of the *n*-1 subjects in those electrodes, (iii) build a regression model of these values against the target variable, and then (iv) make a prediction for the target variable for the left-out subject. This procedure is repeated to obtain an out-of-sample residual (i.e., prediction error) for each subject, and the R^2^ was calculated comparing the variance of the residuals vs the variance of the target variable.

3) Then, we found the optimal threshold ϑ* as the value of ϑ that maximised the out-of-sample R^2^.

4) Finally, we built a final predictor following the same algorithm as in step 2, using all *n* subjects and the optimal correlation threshold ϑ*. The performance of this predictor was assessed by calculating the correlation between predicted values and the target using Spearman’s *r_s_*, and its significance through surrogate data tests (i.e., comparing the out-of-sample R^2^ of the model with the same quantity evaluated on data where the target values have been shuffled between subjects, repeated for 1,000 permutations). These values are reported in the inset of Figs. 4A and 4C.

Please note that it is well known that the method used to choose the threshold— done in this case in step (3)— is known to not affect the false positive rate (i.e. the estimated statistical significance) of the test, but only its power (i.e. its sensitivity).^3^

To test the robustness of the findings obtained via this pipeline, we also investigated the results obtained via a simpler procedure: testing the predictive ability on both insight and well-being changes of simple spatial averages of the LZ values across all of the electrodes at different timepoints (i.e., 1hrs, 2hrs, and 4.5hrs post-dosing). The findings of these additional analyses— with their significance corrected for multiple comparisons via Bonferroni— directly support the findings obtained with the data-driven method described here. These additional, simpler control analyses are reported in the supplementary material, Section 3.4.

**MRI/Neural outcomes.**

To assess enduring brain effects, participants attended three scanning sessions, 1-month apart: scan 1 collected baseline data, scan 2 served as the one-month follow-up for 1 mg control as well as a baseline for post-25mg interventions, and scan 3 served as the one-month follow-up for 25 mg psilocybin and was the primary study endpoint. Imaging was performed on a 3T Siemens Tim Trio using a 12-channel head coil. Whole-head anatomical images were acquired using the Alzheimer’s Disease Neuroimaging Initiative, Grand Opportunity (ADNI-GO)^4^ recommended Magnetization Prepared Rapid Gradient Echo (MPRAGE) parameters – 1 mm isotropic voxels, TR = 2300 ms, TE = 2.98 ms, 160 sagittal slices, 256 × 256 in-plane FOV, flip angle = 9°, bandwidth = 240 Hz/pixel, parallel imaging (PI) factor = 2, Inversion time = 900 ms. The three anatomical images from the different scanning data were merged into one T1 image, so to have better structural resolution. This was done by registering scan 2 and scan 3 to scan 1, and then averaging the scans together.

***Functional MRI procedures.*** T2*-weighted echo-planar images (EPI) were acquired using the MB2R2 protocol^5^ with interleaved slice acquisitions for the functional scans using 3 mm isotropic voxels, TR = 1250 ms, TE = 30 ms, 44 axial slices, 192 mm in-plane FOV, flip angle = 80°, bandwidth = 2232 Hz/pixel, GRAPPA acceleration = 2, number of volumes = 384. Both the resting state and emotional faces paradigm scans were each 8 minutes in duration. Resting-state scans were completed with eyes-closed.

***Pre-processing.***

The preprocessing pipeline used here was similar to our previous studies^6,7^, yet with slight modifications (e.g. not doing slice time correction due to the multiband sequence). FMRIB Software Library (FSL)^8^, Analysis of Functional NeuroImages (AFNI)^9^, Freesurfer^10^ and Advanced Normalization Tools (ANTS)^11^ were used to analyze the resting-state data. Motion was measured using frame-wise displacement (FD)^12^. The criterion for exclusion was subjects with > 20% scrubbed volumes when the scrubbing threshold is FD = 0.4. Two subjects were excluded to high levels of head motion, and 23 subjects were used for the final analysis. The following preprocessing stages were performed: 1) removal of the first three volumes; 2) de-spiking (3dDespike, AFNI); 3) motion correction (3dvolreg, AFNI) by registering each volume to the volume most similar, in the least squares sense, to all others (in-house code); 4) brain extraction (BET, FSL); 5) rigid body registration to anatomical scans (FSL, BBR); 6) non-linear registration to 2mm MNI brain (Symmetric Normalization (SyN), ANTS); 7) scrubbing^13^ using an FD threshold of 0.4 (the mean percentage of volumes scrubbed for scan 1, scan 2, and scan 3 was 0.9 ± 1.6%, 0.9 ± 1.4%, and 2.6 ± 5.3, respectively). Scrubbed volumes were replaced with the mean of the surrounding volumes. Additional preprocessing steps included: 8) spatial smoothing (FWHM) of 6mm (3dBlurInMask, AFNI); 9) band-pass filtering between 0.01 to 0.08 Hz (3dFourier, AFNI); 10) linear and quadratic de-trending (3dDetrend, AFNI); 11) regressing out 9 nuisance regressors (the same bandpass filter was applied on the nuisance regressors): out of these, 6 were motion-related (3 translations, 3 rotations) and 3 were anatomically-related (not smoothed). Specifically, the anatomical nuisance regressors were: 1) ventricles (Freesurfer, eroded in 2mm space), 2) draining veins (DV) (FSL’s CSF minus Freesurfer’s Ventricles, eroded in 1mm space) and 3) local white matter (WM) (FSL’s WM minus Freesurfer’s subcortical grey matter (GM) structures, eroded in 2mm space). Regarding local WM regression^14^, AFNI’s 3dLocalstat was used to calculate the mean local WM time-series for each voxel, using a 25mm radius sphere centered on each voxel.

***Seed-based resting state functional connectivity (RSFC).***

Based on prior hypotheses, four seeds were chosen for these analyses: (i) bilateral parahippocampus (PH), (ii) bilateral amygdala, (iii) ventromedial prefrontal cortex (vmPFC), and (iv) subgenual anterior cingulate cortex (sgACC). The PH seed was constructed by combining the anterior and posterior parahippocampal gyrus from the Harvard-Oxford probabilistic atlas, which was then thresholded at 50%. The vmPFC seed was the same as one previously used by our team in analyses of the acute effects of LSD, psilocybin and MDMA^6^. The sgACC seed was a 5 mm sphere centered at MNI coordinates ±2 28 -5. Bilateral amygdala seed was based on Harvard-Oxford probabilistic atlas, threshold at 50%. Mean time-series were derived for these seeds for each resting-state scan. The RSFC analyses were performed using FSL’s FEAT for each subject. Pre-whitening (FILM) was applied. A higher-level analysis was performed to compare pre-treatment (scan 2) and post-treatment (scan 3) conditions using a mixed-effects GLM (FLAME 1 + 2), cluster corrected (Z>2.3, p<0.05). MRIcron was used to display the results. Only the amygdala results were significant.

***Emotional faces paradigm procedures*.**

The emotional faces paradigm was a block-design task lasting 8 min. Participants used a mirror mounted on the head-coil to view a screen mounted in the rear of the scanner bore, where visual stimuli were back-projected through a wave-guide in the rear wall of the scanner room. Participants were shown faces with either fearful, happy, or neutral expressions, selected from the Karolinska Directed Emotional Faces set^15^. An equal number of male and female faces were selected for the task. Each face was presented on screen for 3 s, and five faces of the same expression were presented in each 15 s block. Rest blocks (also 15 s) were also included, and there were 8 repetitions of each block type, presented in a pseudo-random sequence (32 blocks in total). Three versions of the task were used and the order of the task versions on each scanning visit was counter-balanced across participants. Participants passively viewed the faces but were instructed to press a single button with their thumb with the presentation of each new face, to confirm that they were paying attention to the stimuli.

For the emotional faces paradigm, we used the same resting-state preprocessing pipeline outlined above but with one modification – as in our previous emotional faces research^16^. The scrubbing threshold was increased to 0.9, as this is a threshold that better suits task paradigms^17^. No subjects were excluded due to head motion, and 25 subjects were used for the final analysis. The mean percentage of volumes scrubbed for scan 1, scan 2, and scan 3 was 1.1 ± 1.8%, 1.1 ± 2.3%, and 1.1 ± 1.9, respectively.

Different approaches were used to investigate changes in amygdala response in this paradigm: (i) voxelwise analysis within a bilateral amygdala mask (Harvard-Oxford atlas, probability >50%); (ii) calculating mean amygdala signal of left and right amygdala ROIs; and (iii) A whole-brain voxelwise analysis. For all approaches, a standard GLM was used for the first analysis step, as implemented in the FEAT module in FSL. Regressors derived from the onset times of each stimulus condition were convolved with a Gamma function in order to simulate the Haemodynamic Response Function (HRF). Prewhitening (FILM) was applied to correct for autocorrelations. Contrasts were defined that isolated activity related to each stimulus condition (fearful, happy, neutral) relative to the baseline, and comparisons were also made that contrasted between stimulus conditions, as appropriate i.e., [ fearful > happy+neutral ] and [ happy > fearful+neutral ]. Mixed-effects GLM (FLAME-1+2) was used for the voxelwise analysis with a statistical threshold of Z>2.3, (cluster-corrected for multiple comparisons, p<0.050).

***Generalized psychophysiological interaction***

To identify condition-specific modulation of amygdala functional connectivity (FC), we utilized a generalized psychophysiological (gPPI) approach using the CONN fMRI toolbox (https://www.nitrc.org/projects/ conn). Preprocessing steps included anatomical component correction (aCompCor) to remove noise and movement confounds on a voxel-to-voxel basis; denoising with parameters for white matter (5P), cerebrospinal fluid (5P), realignment (6P) and scrubbed volumes; bandpass filtering (0.008-0.09 Hz), and linear detrending after regression. The gPPI was computed using bivariate regression in CONN by specifying the following regressors: post-1mg fear, post-1mg happy, post-1mg neutral, post-25mg fear, post-25mg happy, post-25mg neutral, time-series regressor for the seed region (left and right amygdala), and PPI terms for the fear, happy, and neutral conditions. To minimize the number of statistical tests, only significant condition contrasts from the BOLD analysis were entered into analyses. Left and right amygdala seeds were run separately with alpha set at a false discovery rate (FDR)-corrected ‘q’ value less than 0.05. FDR correction (*q*_FDR_ = 0.05) was also used to adjust for laterality (left and right amygdala seeds), and for the 3 condition contrasts of interest (fear, fear > happy + neutral, and happy > fear + neutral).

***Brain network modularity.***

Brain network modularity was computed on cortical (200 x 200)^18^ interregional RSFC. As summarized by the Q value (see Star Methods), modularity is a measure of the decomposability of brain connectivity into distinct modules, where each module represents a set of brain regions that exhibit strong RSFC with each other (intra-modular RSFC) and weaker RSFC with regions of other modules (inter-modular RSFC).

For the modularity analyses, the RS-fMRI timeseries data was parcellated into 200 regions based on the Schaefer local-global parcellation^18^ and the Pearson’s (r) correlation was computed between regional timeseries to create a 200 x 200 RSFC matrix. Modularity was then computed on this (unthresholded) matrix using the Louvain algorithm^19^ implemented with the Network Community Toolbox (<http://commdetect.weebly.com/>), treating negative values asymmetrically. This algorithm finds modular partitions of the network (i.e., RSFC matrix) which optimize the modularity value, Q, by grouping nodes into non-overlapping modules (sets of regions) that maximize intra-modular and minimize inter-modular connections^20^. The modularity value Q for a given modular partition is computed as follows:

$$Q= \frac{1}{l}\sum_{i,j\in N} \left[ w_{ij}-\frac{k_{i}k_{j}}{l} \right]\delta_{m_{i}mj}$$

where 𝑤 is the edge weight (i.e., functional connectivity value) between nodes *i* and *j*, 𝑙^w^ is the sum of all weights in the graph, 𝑘_i_ is the weighted degree (edge weight summed across all edges) of node *i*, and 𝑚_i_ is a module containing node i. $\delta_{m_{i}mj}$ =1 if nodes *i* and *j* belong to the same module, and = 0 otherwise. The Q value for a given partition therefore quantifies the strength of within-module edges relative to the strength of between-module edges, or, in other words, the extent to which distinct modules can be delineated in the data. This algorithm has a single free parameter, gamma (γ), which controls how many modules will be detected. We kept this at the default value of 1. The Louvain algorithm was run iteratively 100 times at the individual-subject level and the average Q value across these runs was used for each subject. Given that modularity is sensitive to the average correlation strength of the network, Q values were normalized for each subject by the mean Q value generated by 100 randomly shuffled (‘rewired’) permutations of that subject’s RSFC matrix.

***Structural MRI procedures.*** Diffusion-weighted MRI (dMRI) data was acquired with the following acquisition parameters: 64 directions with b = 1000 s/mm^2^; 6 images without diffusion-weighting; 1 image without diffusion-weighting and with the phase encoding direction reversed; TE = 88 ms; TR = 3010 ms; voxel size = 1.9 x 1.9 x 2.0 mm^3^; 72 slices. The acquisition was repeated three times to maximize signal-to-noise ratio (SNR). The images were preprocessed using MRtrix3^21^ and the following steps: random matrix denoising^22,23^, Gibbs-ringing reduction^24^, distortion and motion correction^8,25^, and bias field correction^26^.

For distortion correction, MRtrix's `dwifslpreproc` was used to call FSL's `eddy` and `topup`. `eddy` was used to correct eddy current-induced distortions and subject movements, while `topup` was used to correct susceptibility-induced distortions.

Region of interest (ROI) analysis was chosen for studying white matter plasticity. TractSeg’s pre-trained neural network^27^ was used to segment white matter and its outputs, representing probabilities of the tract being present in a voxel, were converted into binary masks with a threshold of 0.975 for reliable and reproducible segmentation. The following association fibers were included in the analysis: arcuate fascicle, cingulum, inferior occipito-frontal fascicle, inferior longitudinal fascicle, middle longitudinal fascicle, superior longitudinal fascicle, and uncinate fascicle. Furthermore, the following were included to incorporate the major frontal lobe tracts: anterior thalamic radiation, rostrum of the corpus callosum, genu of the corpus callosum, striato-prefontal tract (PFC-STR), and thalamo-prefrontal tract (PFC-THA). ROIs from the different hemispheres were merged.

The diffusion tensor was estimated in each voxel using an iterated weighted least squares algorithm in Mrtrix3^21,28,29^. The mean values of axial diffusivity (AD) and radial diffusivity (RD) were calculated in each ROI for each subject and time point. Statistical analysis was performed on AD and RD that measure diffusion along orthogonal directions and are independent. In addition, diffusion tensors were estimated with a free-water component, done to remove partial volume effects^30^. Repeated measures ANOVA was performed on AD and RD separately for each tract with Bonferroni multiple comparisons correction. The analysis was then repeated for mean diffusivity (MD) and fractional anisotropy (FA).

A study-specific template was generated, which was used as the registration target in a Tract-Based Spatial Statistics (TBSS) pipeline ^31^. A white matter skeleton was created from the averaged and registered FA images, which was thresholded at FA < 0.2. DWI metrics were then projected onto the skeleton to perform statistical analysis using FSL randomise.

**Psychological outcomes**

***Cognitive flexibility.*** Cognitive flexibility was measured via an intra-dimensional/extra-dimensional (IDED) task that had been optimised for internet-based delivery^32,33^. The IDED task consists of 9 stages that require simple discrimination learning, compound discrimination learning, abstraction, attentional set-shifting, and reversal learning (see supplementary material for more detail). Two stimuli are presented to the participant at a time. At any given phase, a hidden rule determines the correct response to the trial. Participants are required to determine the rule and chose the correct stimuli in 6 consecutive trials before progressing to the next phase. Participants are not informed when the rule changes, only receiving ticks or crosses as feedback for correct or incorrect responses, respectively. Participants fail the task upon responding incorrectly 30 times within any given phase. Stimuli sets are pseudorandomized (different sets of lines and shapes from a given pool) between timepoints. Errors made at each phase were adjusted to the total number of stimuli presented within each phase, representing a measure of ‘phase-accuracy’ in order to assess changes in IDED performance across time-points while controlling for individual variability in learning rates. This adjustment was made by dividing the number of errors made by the number of trials completed per phase, therefore providing a representation of the percentage of trials in which an error was made. Participants completed the IDED task 1-day before and one-month after each dosing day.

***Insight.*** Psychological insight was measured via the Psychological Insight Scale (PIS)^34^. The PIS is a 6-7 item questionnaire that measures psychological insightfulness following a psychedelic experience (PIS-6) and accompanied behavioural changes (PIS-7). The PIS is scored using a VAS (0–100, with incremental units of one) with zero defined as ‘no more than usual’ and 100 defined as ‘much more than usual’. Participants completed the PIS at 1-day, 2-weeks and one-month after each dosing day.

***Well-being.*** Psychological well-being was measured using the 14-item *Well-being Warwick-Edinburgh Mental Wellbeing Scale* (WEMWBS)^35^. The WEMWBS is designed to assess mental well-being itself and not the determinants of mental well-being – e.g. resilience, problem solving, etc. The WEMWBS includes hedonic (i.e. happiness, life satisfaction) and eudaimonic (i.e. positive relationships, psychological functioning) items which together measure mental well-being. Items are rated on a 5-point Likert scale (from 1 = none of the time to 5 = all of the time) to yield a total summed score, with a minimum possible score of 14 and a maximum score of 70 (population norms: *M*=51, *SD*=9)^36^. Participants completed the WEMWBS 1-day before and at 2-weeks and 4-weeks after each dosing day.

***Retrospective ranking of the experience***. Broadly inspired by (but not directly related to) previous methods by Griffiths et al^37^, six months post-dosing, participants reflected on the peak of each dosing session and answered the following three questions: 1) “How profound was the state of consciousness you experienced?”, 2) “How intense was the state of consciousness you experienced?”, and 3) “How unusual was the state of consciousness you experienced?” (See Figure S26). Each item was rated using an eight-point Likert scale, from 1 = “no more ‘profound / intense / unusual’ than routine everyday states of consciousness” to 8 = “the single most ‘profound / intense / unusual’ state of consciousness of my life, that I can recall”).

**QUANTIFICATION AND STATISTICAL ANALYSIS**

All statistical analyses were performed using either SPSS version 26.0 (IBM Corp., Armonk, NY, USA), R Studio 2022.07.1 (RStudio PBC, Boston,. MA, USA), SciPy 1.9.1 and statsmodels 0.13.2 in Python 3.9.13 (Python Software Foundation), or MATLAB 2019b (MathWorks, Natick, USA) and illustrated using GraphPad Prism version 9.2 (GraphPad Software, La Jolla California USA). Data were subjected to one- and two-way repeated measures ANOVAs or linear mixed-effects (LME) model, where appropriate, and followed up on with *post-hoc* tests corrected for multiple comparisons. The structure of random effects in LME models was built following standard recommendations^38^. The 95% CIs are provided. Effect sizes were calculated using the repeated measures Cohen’s *d* formula. Statistical significance was assumed at the p≤0.050 probability level.

1	Oostenveld, R., Fries, P., Maris, E. & Schoffelen, J. M. FieldTrip: Open source software for advanced analysis of MEG, EEG, and invasive electrophysiological data. *Comput Intell Neurosci* **2011**, 156869 (2011). <https://doi.org/10.1155/2011/156869>

2	Madsen, M. K. *et al.* Psilocybin-induced changes in brain network integrity and segregation correlate with plasma psilocin level and psychedelic experience. *Eur Neuropsychopharmacol* **50**, 121-132 (2021). <https://doi.org/10.1016/j.euroneuro.2021.06.001>

3	Maris, E. & Oostenveld, R. Nonparametric statistical testing of EEG- and MEG-data. *J Neurosci Methods* **164**, 177-190 (2007). <https://doi.org/10.1016/j.jneumeth.2007.03.024>

4	Jack, C. R., Jr. *et al.* The Alzheimer's Disease Neuroimaging Initiative (ADNI): MRI methods. *J Magn Reson Imaging* **27**, 685-691 (2008). <https://doi.org/10.1002/jmri.21049>

5	Demetriou, L. *et al.* A comprehensive evaluation of increasing temporal resolution with multiband-accelerated protocols and effects on statistical outcome measures in fMRI. *NeuroImage* **176**, 404-416 (2018). <https://doi.org/10.1016/j.neuroimage.2018.05.011>

6	Carhart-Harris, R. L. *et al.* Neural correlates of the LSD experience revealed by multimodal neuroimaging. *Proceedings of the National Academy of Sciences of the United States of America* **113**, 4853-4858 (2016). <https://doi.org/10.1073/pnas.1518377113>

7	Carhart-Harris, R. L. *et al.* Psilocybin for treatment-resistant depression: fMRI-measured brain mechanisms. *Scientific Reports* **7**, 13187 (2017). <https://doi.org/10.1038/s41598-017-13282-7>

8	Smith, S. M. *et al.* Advances in functional and structural MR image analysis and implementation as FSL. *NeuroImage* **23 Suppl 1**, S208-219 (2004). <https://doi.org/10.1016/j.neuroimage.2004.07.051>

9	Cox, R. W. AFNI: software for analysis and visualization of functional magnetic resonance neuroimages. *Comput Biomed Res* **29**, 162-173 (1996). <https://doi.org/10.1006/cbmr.1996.0014>

10	Dale, A. M., Fischl, B. & Sereno, M. I. Cortical surface-based analysis. I. Segmentation and surface reconstruction. *NeuroImage* **9**, 179-194 (1999). <https://doi.org/10.1006/nimg.1998.0395>

11	Avants, B. B., Tustison, N. & Song, G. Advanced normalization tools (ANTS). *Insight j* **2**, 1-35 (2009).

12	Power, J. D. *et al.* Methods to detect, characterize, and remove motion artifact in resting state fMRI. *NeuroImage* **84**, 320-341 (2014). <https://doi.org/10.1016/j.neuroimage.2013.08.048>

13	Power, J. D., Barnes, K. A., Snyder, A. Z., Schlaggar, B. L. & Petersen, S. E. Spurious but systematic correlations in functional connectivity MRI networks arise from subject motion. *NeuroImage* **59**, 2142-2154 (2012). <https://doi.org/10.1016/j.neuroimage.2011.10.018>

14	Jo, H. J., Saad, Z. S., Simmons, W. K., Milbury, L. A. & Cox, R. W. Mapping sources of correlation in resting state FMRI, with artifact detection and removal. *NeuroImage* **52**, 571-582 (2010). <https://doi.org/10.1016/j.neuroimage.2010.04.246>

15	Goeleven, E., De Raedt, R., Leyman, L. & Verschuere, B. Vol. 22	1094-1118 (Taylor & Francis, United Kingdom, 2008).

16	Roseman, L., Demetriou, L., Wall, M. B., Nutt, D. J. & Carhart-Harris, R. L. Increased amygdala responses to emotional faces after psilocybin for treatment-resistant depression. *Neuropharmacology* **142**, 263-269 (2018). <https://doi.org/10.1016/j.neuropharm.2017.12.041>

17	Siegel, J. S. *et al.* Statistical improvements in functional magnetic resonance imaging analyses produced by censoring high-motion data points. *Hum Brain Mapp* **35**, 1981-1996 (2014). <https://doi.org/10.1002/hbm.22307>

18	Schaefer, A. *et al.* Local-Global Parcellation of the Human Cerebral Cortex from Intrinsic Functional Connectivity MRI. *Cerebral Cortex* **28**, 3095-3114 (2017). <https://doi.org/10.1093/cercor/bhx179>

19	Blondel, V. D., Guillaume, J.-L., Lambiotte, R. & Lefebvre, E. Fast unfolding of communities in large networks. *Journal of statistical mechanics: theory and experiment* **2008**, P10008 (2008).

20	Newman, M. E. Fast algorithm for detecting community structure in networks. *Physical review E* **69**, 066133 (2004).

21	Tournier, J. D. *et al.* MRtrix3: A fast, flexible and open software framework for medical image processing and visualisation. *NeuroImage* **202**, 116137 (2019). <https://doi.org/10.1016/j.neuroimage.2019.116137>

22	Cordero-Grande, L., Christiaens, D., Hutter, J., Price, A. N. & Hajnal, J. V. Complex diffusion-weighted image estimation via matrix recovery under general noise models. *NeuroImage* **200**, 391-404 (2019). <https://doi.org/10.1016/j.neuroimage.2019.06.039>

23	Veraart, J. *et al.* Denoising of diffusion MRI using random matrix theory. *NeuroImage* **142**, 394-406 (2016). <https://doi.org/10.1016/j.neuroimage.2016.08.016>

24	Kellner, E., Dhital, B., Kiselev, V. G. & Reisert, M. Gibbs-ringing artifact removal based on local subvoxel-shifts. *Magn Reson Med* **76**, 1574-1581 (2016). <https://doi.org/10.1002/mrm.26054>

25	Andersson, J. L. R. & Sotiropoulos, S. N. An integrated approach to correction for off-resonance effects and subject movement in diffusion MR imaging. *NeuroImage* **125**, 1063-1078 (2016). <https://doi.org/10.1016/j.neuroimage.2015.10.019>

26	Tustison, N. J. *et al.* N4ITK: improved N3 bias correction. *IEEE Trans Med Imaging* **29**, 1310-1320 (2010). <https://doi.org/10.1109/tmi.2010.2046908>

27	Wasserthal, J., Neher, P. & Maier-Hein, K. H. TractSeg - Fast and accurate white matter tract segmentation. *NeuroImage* **183**, 239-253 (2018). <https://doi.org/10.1016/j.neuroimage.2018.07.070>

28	Basser, P. J., Mattiello, J. & LeBihan, D. MR diffusion tensor spectroscopy and imaging. *Biophysical journal* **66**, 259-267 (1994). <https://doi.org/10.1016/s0006-3495(94)80775-1>

29	Veraart, J., Sijbers, J., Sunaert, S., Leemans, A. & Jeurissen, B. Weighted linear least squares estimation of diffusion MRI parameters: strengths, limitations, and pitfalls. *NeuroImage* **81**, 335-346 (2013). <https://doi.org/10.1016/j.neuroimage.2013.05.028>

30	Pasternak, O., Sochen, N., Gur, Y., Intrator, N. & Assaf, Y. Free water elimination and mapping from diffusion MRI. *Magnetic Resonance in Medicine* **62**, 717-730 (2009). <https://doi.org/https://doi.org/10.1002/mrm.22055>

31	Smith, S. M. *et al.* Tract-based spatial statistics: voxelwise analysis of multi-subject diffusion data. *Neuroimage* **31**, 1487-1505 (2006). <https://doi.org/10.1016/j.neuroimage.2006.02.024>

32	Roberts, A. C., Robbins, T. W. & Everitt, B. J. The effects of intradimensional and extradimensional shifts on visual discrimination learning in humans and non-human primates. *Q J Exp Psychol B* **40**, 321-341 (1988).

33	Owen, A. M., Roberts, A. C., Polkey, C. E., Sahakian, B. J. & Robbins, T. W. Extra-dimensional versus intra-dimensional set shifting performance following frontal lobe excisions, temporal lobe excisions or amygdalo-hippocampectomy in man. *Neuropsychologia* **29**, 993-1006 (1991). <https://doi.org/10.1016/0028-3932(91)90063-e>

34	Peill, J. M. *et al.* Validation of the Psychological Insight Scale: A new scale to assess psychological insight following a psychedelic experience. *Journal of psychopharmacology (Oxford, England)* **36**, 31-45 (2022). <https://doi.org/10.1177/02698811211066709>

35	Tennant, R. *et al.* The Warwick-Edinburgh Mental Well-being Scale (WEMWBS): development and UK validation. *Health and Quality of Life Outcomes* **5**, 63 (2007). <https://doi.org/10.1186/1477-7525-5-63>

36	Morris, S. & Earl, K. Health Survey for England 2016: Well-being and mental health. *Leeds, UK: Health and Social Care Information Centre. Retrieved from* [*http://healthsurvey.hscic.gov.uk/media/63763/HSE2016-Adult-wel-bei.pdf*](http://healthsurvey.hscic.gov.uk/media/63763/HSE2016-Adult-wel-bei.pdf) (2017).

37	Griffiths, R. R., Richards, W. A., McCann, U. & Jesse, R. Psilocybin can occasion mystical-type experiences having substantial and sustained personal meaning and spiritual significance. *Psychopharmacology* **187**, 268-283; discussion 284-292 (2006). <https://doi.org/10.1007/s00213-006-0457-5>

38	Barr, D. J., Levy, R., Scheepers, C. & Tily, H. J. Random effects structure for confirmatory hypothesis testing: Keep it maximal. *J Mem Lang* **68**, 10.1016/j.jml.2012.1011.1001 (2013). <https://doi.org/10.1016/j.jml.2012.11.001>
